# Interpreting *cis*-regulatory mechanisms from genomic deep neural networks using surrogate models

**DOI:** 10.1101/2023.11.14.567120

**Authors:** Evan E Seitz, David M McCandlish, Justin B Kinney, Peter K Koo

**Affiliations:** Simons Center for Quantitative Biology, Cold Spring Harbor Laboratory, Cold Spring Harbor, NY, USA

## Abstract

Deep neural networks (DNNs) have greatly advanced the ability to predict genome function from sequence. Interpreting genomic DNNs in terms of biological mechanisms, however, remains difficult. Here we introduce SQUID, a genomic DNN interpretability framework based on surrogate modeling. SQUID approximates genomic DNNs in user-specified regions of sequence space using surrogate models, i.e., simpler models that are mechanistically interpretable. Importantly, SQUID removes the confounding effects that nonlinearities and heteroscedastic noise in functional genomics data can have on model interpretation. Benchmarking analysis on multiple genomic DNNs shows that SQUID, when compared to established interpretability methods, identifies motifs that are more consistent across genomic loci and yields improved single-nucleotide variant-effect predictions. SQUID also supports surrogate models that quantify epistatic interactions within and between *cis*-regulatory elements. SQUID thus advances the ability to mechanistically interpret genomic DNNs.

---

Deep neural networks (DNNs) are increasingly being used to analyze and biologically interpret functional genomics data. DNNs have demonstrated remarkable success at predicting diverse genomic activities from primary genome sequences, including mRNA expression levels,^1^ transcription start site activity,^2, 3^ mRNA splicing patterns,^4^ protein-DNA binding,^5^ chromatin accessibility,^6^ and chromatin conformation.^7^ It is widely believed that the success of genomic DNNs reflects their ability to accurately model complex biological mechanisms. However, interpreting genomic DNNs in terms of biological mechanisms remains difficult.

A variety of *post hoc* attribution methods have been developed for aiding the interpretation of genomic DNNs.^8, 9^ The most commonly used attribution methods produce *attribution maps*, which quantify the position-specific effects that variant nucleotides in a sequence of interest have on DNN predictions. Attribution maps are often mechanistically interpreted by identifying motifs, i.e. recurrent sequence patterns characteristic of specific biological mechanisms, present within these maps. To provide consistent biological explanations for sequence activity, attribution methods must produce consistent motifs across input sequences that function through shared biological mechanisms.

Established attribution methods for genomic DNNs have major limitations. Different attribution methods use different strategies to quantify position-specific nucleotide effects, and can therefore yield different mechanistic explanations.^10–12^ For example, Saliency Maps^13^ quantify nucleotide effects using the gradient of DNN predictions at the sequence of interest, whereas DeepLIFT^14^ propagates activation differences between the sequence of interest and a reference sequence. Moreover, the most widely-used attribution methods in genomics – including Saliency Maps,^13^ DeepLIFT,^14^ *in silico* mutagenesis (ISM),^15^ SmoothGrad,^16^ Integrated Gradients,^17^ and DeepSHAP^18^ – assume that nucleotide effects on DNN predictions are locally additive. As a result, these attribution methods do not explicitly account for the combinations of genetic interactions (i.e., specific epistasis^19–22^), global nonlinearities (i.e., global epistasis^23–27^), and heteroscedastic noise^26^ that are often present in functional genomics data.

Here we introduce SQUID (**S**urrogate **Qu**antitative **I**nterpretability for **D**eepnets), an interpretability framework for genomic DNNs that overcomes these limitations. SQUID uses surrogate models—simple models with interpretable parameters—to approximate the DNN function within localized regions of sequence space. SQUID applies MAVE-NN,^26^ a quantitative modeling framework developed for analyzing multiplex assays of variant effects (MAVEs), to *in silico* MAVE datasets generated using the DNN as an oracle. SQUID models DNN predictions in a user-specified region of sequence space; this is similar to LIME^28^, a surrogate modeling approach developed for general DNNs. Unlike LIME, however, SQUID is able to account for important domain-specific outputs of genomic DNNs: global nonlinearities and heteroskedastic noise. Indeed, SQUID can be viewed as a generalization of LIME; see Methods for details.

Benchmarking SQUID against existing attribution methods, we find that SQUID more consistently quantifies the binding motifs of transcription factors (TFs), reduces noise in attribution maps, and improves variant-effect predictions. We also find that the domain-specific surrogate models used by SQUID are critical for this improved performance. Finally, we show how SQUID can provide insights into epistatic interactions in *cis*-regulatory elements, and can be used to study such interactions both locally and globally in sequence space. SQUID thus provides a new and useful way to interpret genomic DNNs.

## Results

### SQUID: An interpretable surrogate modeling framework for genomic DNNs

SQUID approximates DNNs in user-specified regions of sequence space using surrogate models that have mechanistically interpretable parameters. The SQUID framework comprises three steps (Fig.1a): (1) generate an *in silico* MAVE dataset comprised of variant sequences and the corresponding DNN predictions; (2) fit a surrogate model to these *in silico* data using MAVE-NN^26^; and (3) visualize and interpret the surrogate model’s parameters. This workflow requires the specification of two key analysis parameters: the region of sequence space over which the DNN is to be approximated, and the mathematical form of the surrogate model. By choosing different values for these analysis parameters, users are able to test different mechanistic hypotheses. SQUID assists users by facilitating the use of analysis parameters that have been found in practice to work well in the design and analysis of MAVE experiments.

First, SQUID generates an *in silico* MAVE dataset. This is done by generating a library of variant sequences, then using the DNN to assign a functional score to each sequence in the library. In this paper we consider two types of libraries: local and global. A local library is generated by partially mutagenizing a specific sequence of interest (e.g., a genomic *cis*-regulatory sequence). A global library is generated by inserting partially-mutagenized versions of a genetic element of interest into random sequences. In what follows, we use a partial mutagenesis rate of 10% per nucleotide (which is common in MAVE experiments, e.g. ref.^29^) and libraries comprising 100,000 variant sequences (unless otherwise noted).

Next, SQUID fits a surrogate model to the *in silico* MAVE dataset. SQUID uses nonlinear surrogate models developed specifically for modeling MAVE data.^26^ These models, called latent phenotype models, have three components (Fig. 1b): a genotype-phenotype (G-P) map, a global epistasis (GE) nonlinearity, and a noise model. The G-P map projects the input sequence down to a one-dimensional latent phenotype. SQUID, via MAVE-NN, supports the use of additive G-P maps, pairwise-interaction G-P maps, and user-defined G-P maps. The GE nonlinearity, which is modeled using a linear combination of sigmoids, maps the latent phenotype to a most-probable DNN prediction. The noise model then describes how actual DNN predictions are expected to scatter around the predicted most-probable value. SQUID supports a variety of noise models, including heteroscedastic noise models based on the skewed-t distribution of Jones and Faddy^30^ (which we use in this paper). Surrogate model parameters are inferred from *in silico* MAVE data by maximizing variational information^26^ (or equivalently, log likelihood) using standard stochastic gradient optimization.

**Figure 1.**
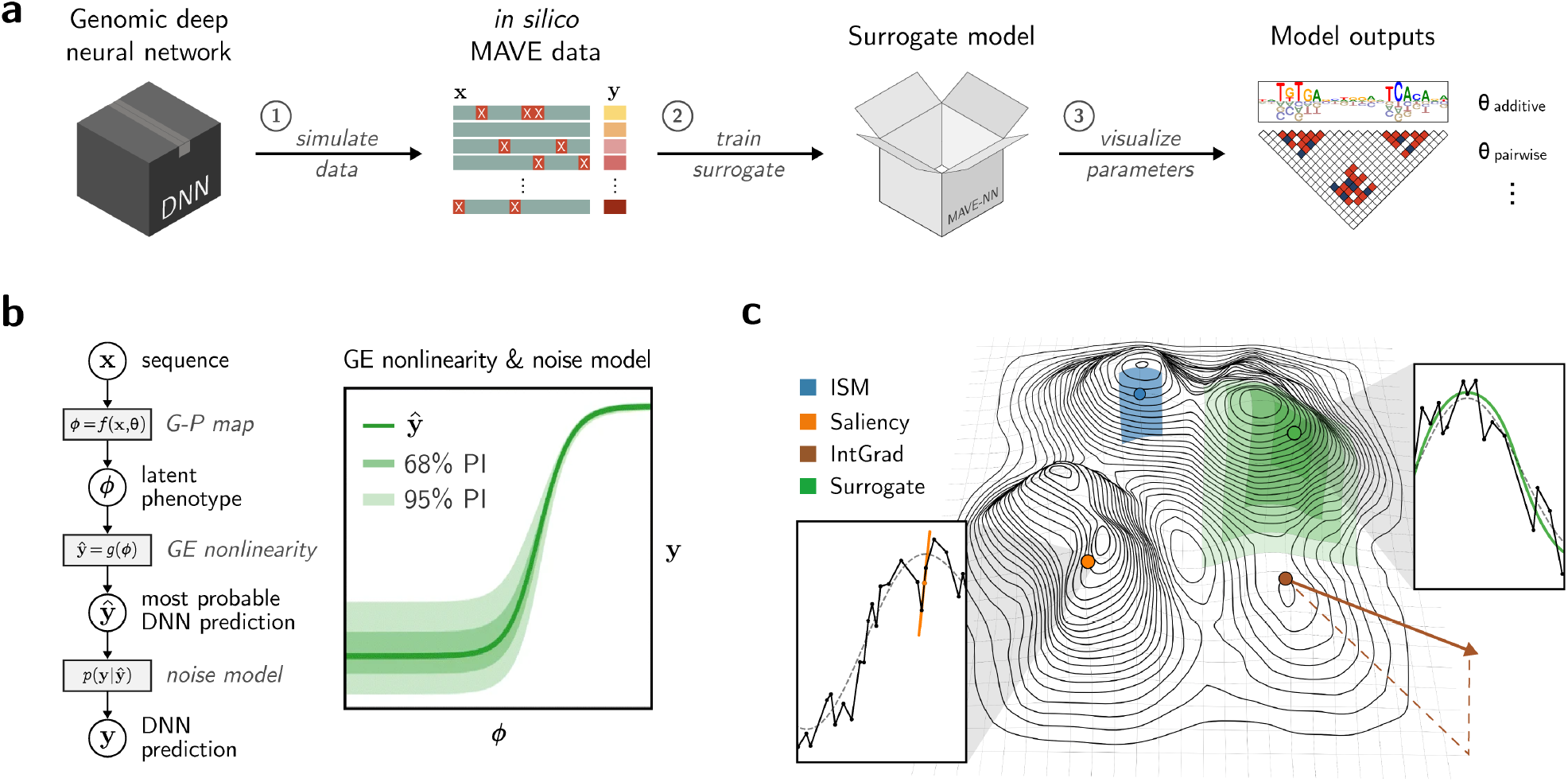
Overview of SQUID. **a**, Schematic of the SQUID modeling framework. Analysis using SQUID comprises three main steps: (1) generate an *in silico* MAVE dataset; (2) train a surrogate model on the MAVE dataset; and (3) visualize parameters of the surrogate model to uncover biological mechanisms. **b**, Structure of the latent phenotype surrogate models supported by SQUID. G-P, genotype-phenotype; GE, global epistasis; PI, prediction interval. **c**, Schematic diagram of a DNN function on a 2-dimensional projection of sequence space. Each point in the plane corresponds to a unique sequence, and elevations represent DNN predictions. Green region schematizes the ability of surrogate models to approximate the DNN function over an extended region of sequence space. Insets show an example DNN function (in 1D profile; black line) centered about a sequence of interest with the ground-truth function (dashed line) overlaid. Left inset illustrates the sensitivity of Saliency Maps to non-smooth local function properties. Right inset illustrates the ability of surrogate models to better approximate ground truth. ISM, *in silico* mutagenesis; IntGrad, Integrated Gradients.

The parameters of the G-P map are of primary interest for downstream mechanistic interpretations. The parameters of additive G-P maps quantify single-nucleotide effects, and are readily visualized using the same methods (sequence logos^31^ and heatmaps) normally used for standard attribution maps. The parameters of pairwise-interaction G-P maps quantify epistatic genetic interactions as well as single-nucleotide effects, and can be visualized using block-triangular heat maps (as in ref.^26^).

### SQUID improves the quantification of transcription factor binding motifs

A common goal when interpreting attribution maps for a DNA sequence of interest is to identify functional binding sites for TFs. However, the TF binding motifs observed in attribution maps often vary substantially from sequence to sequence. Some of this variation is likely due to underlying biology; for example, TF binding preferences have been experimentally observed to change in response to the binding of other TFs nearby^32–34^. However, variation in TF binding motifs can be exacerbated by the specific ways in which attribution methods quantify the behavior of the DNN function in localized regions of sequence space (Fig. 1c). The consistency of binding motifs observed in attribution maps across different genomic sequences therefore provides a way to quantify and compare the performance of different attribution methods.

To benchmark the consistency of binding motifs identified by different attribution methods, we identified sequences in the human genome that contain putative TF binding sites. Specifically, for each TF, we identified genomic instances of the consensus TF binding site sequence at which a strong motif was observed according to a baseline attribution method. We then aligned these genomic sequences about their putative binding sites and computed, for each genomic sequence, an attribution map that spans the core putative binding site as well as a specified amount of flanking DNA on either side. Each attribution map was then normalized to control for locus-to-locus variation in overall motif scale (scale was substantial). Finally, we calculated the Euclidean distance between the vector of normalized attribution scores for individual sequences and the mean vector of normalized attribution scores. We refer to this distance as the “attribution variation” (Fig. 2a; see Methods for details).

**Figure 2.**
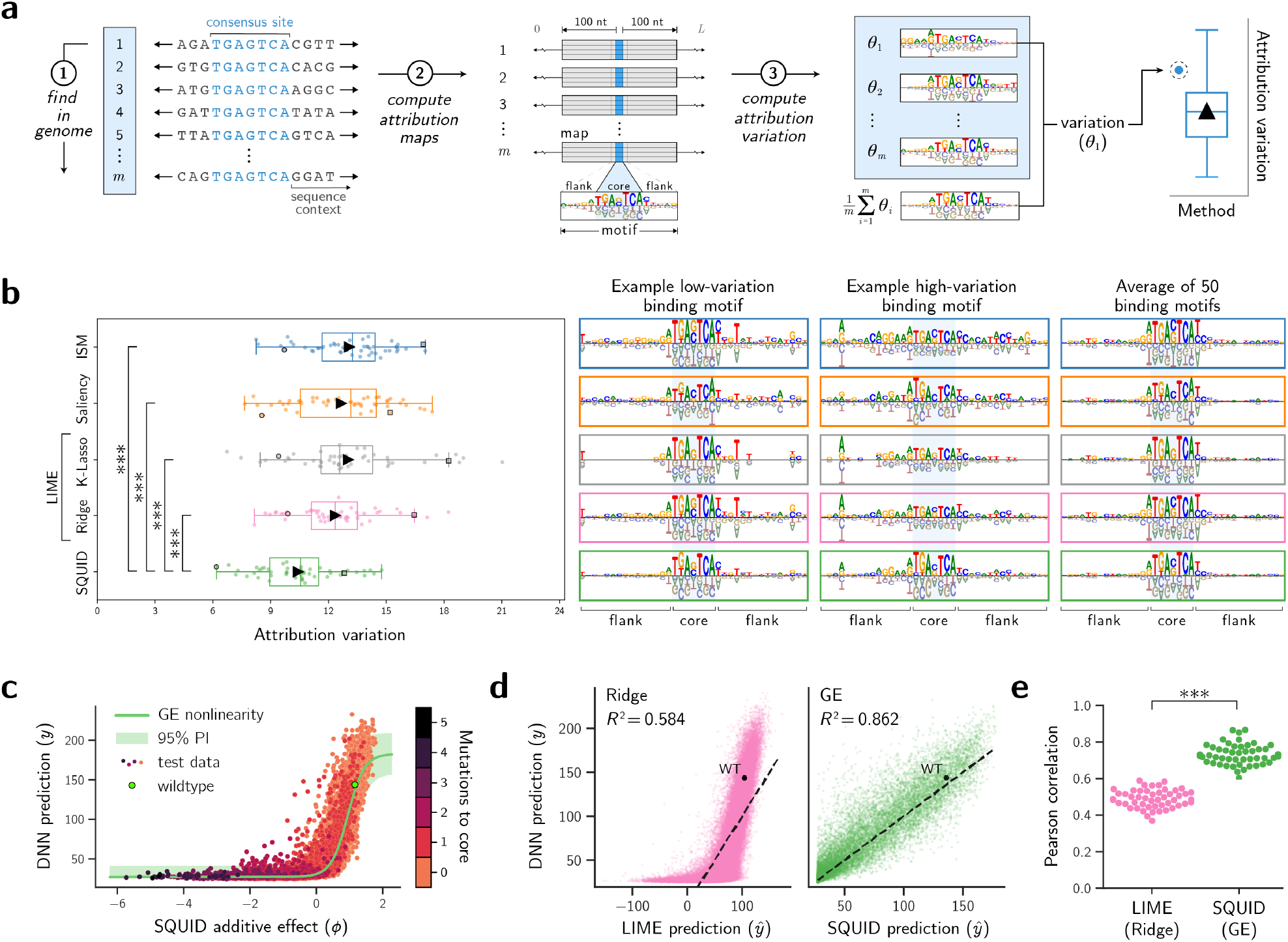
Benchmark analysis of attribution methods. **a**, Our benchmark analysis pipeline consisted of 3 steps: (1) genomic sequences that contained consensus binding sites for a TF of interest were identified; (2) an attribution map spanning the core identified site and 100 nt of flanking sequence on each side was computed for each identified sequence; and (3) a corresponding attribution variation was computed (see Methods for details). **b**, Attribution maps and attribution variation for 7-nt consensus AP-1 binding sites (core sequence `TGAGTGA`; flank size 15 nt), computed using ResidualBind-32 as the DNN. *Left*, Box plots show attribution variation for 50 genomic sequences. Triangles represent means; lines represent median, upper quartile, and lower quartile; whiskers represent 1.5x the inter-quartile range. *Right*, Binding motifs observed for two example genomic sequences, together with the ensemble-averaged binding motif. Note: we used *K* = 50 for the K-Lasso implementation of LIME; results for other choices of *K* are presented in Extended Data Figure 1. **c**, Mutational effects predicted by the DNN versus additive effects predicted by SQUID. Dots represent effects of different sets of mutations to a representative genomic sequence. The GE nonlinearity and 95% PI inferred by SQUID are shown for comparison. **d**, DNN predictions versus the predictions of two surrogate models for a representative genomic sequence. One surrogate model (GE, right) has a GE nonlinearity, the other (Ridge, left) does not. Dots represent test sequences from the *in silico* MAVE dataset. Diagonal lines represent equality between DNN and surrogate model predictions. WT, wild-type sequence; *R*^2^, squared Pearson correlation coefficient. **e**, *R*^2^ values computed as in panel **d** for 50 different sequences of interest. *p*-values in panels **b** and **e** were computed using a one-sided Mann-Whitney U test; ***, *p* < 0.001. TF, transcription factor; DNN, deep neural network; GE, global epistasis; PI, prediction interval; ISM, *in silico* mutagenesis.

We first applied this benchmarking pipeline to the human TF AP-1 using ResidualBind-32,^35^ a genomic DNN that predicts chromatin accessibility in human cell lines. We compared SQUID to two commonly used attribution methods that had previously been used in ref.^35^ to analyze ResidualBind-32: *in silico* mutagenesis (ISM) and Saliency Maps. We also compared SQUID to two different implementations of LIME. One implementation, Ridge, similar to SQUID but lacks a GE nonlinearity and uses a homoscedastic Gaussian noise model. The other implementation, K-Lasso, uses the specific algorithm described in ref.^28^, and requires users to specify the number of nonzero features in the attribution map.

We found that, across different genomic sequences that contain the consensus AP-1 binding site `TGAGTCA`, SQUID recovers AP-1 binding motifs that have markedly-lower attribution variation than the binding motifs recovered by ISM, Saliency Maps, and both implementations of LIME (Fig. 2b, left). This result is supported by the examination of attribution maps of individual sequences (Fig. 2b, right). Compared to the attribution maps provided by other methods, the attribution maps provided by SQUID exhibit core regions with greater similarity to the ensemble-averaged motif, and flanking regions with reduced (likely non-biological) scores. Visual examination of individual PA-1 motifs suggest that some of the observed motif variation was biological in nature, and that in such cases SQUID was better able than other methods to discern functional variation from noise in these cases. Moreover, the motifs provided by SQUID were largely robust to the choice of SQUID analysis parameters (Extended Data Fig. 2). We conclude that SQUID quantifies the AP-1 binding motif from ResidualBind-32 more consistently than does ISM, Saliency Maps, or LIME.

We next investigated whether the surrogate models for AP-1 identified by SQUID benefited from incorporating a GE nonlinearity. We found that SQUID quantifies the AP-1 binding motif better when using GE regression than when using the Ridge implementation of LIME (see Fig. 2b). Plotting the effects that mutations in a representative genomic sequence have on DNN predictions, we found that virtually all combinations of 2 or more mutations to the core 7-nt AP-1 site reduced DNN predictions to near-background levels (Fig. 2c). Moreover, the GE nonlinearity learned by SQUID as part of the surrogate model accurately recapitulated this saturation effect. By contrast, surrogate modeling using the Ridge implementation of LIME failed to capture this saturation effect (Figs. 2d and 2e). This finding demonstrates that surrogate modeling of genomic DNNs can benefit from using our domain-specific surrogate models, as opposed to the linear surrogate models that are standard in other fields (e.g., computer vision^28^).

We then expanded our analysis to other TFs and to other genomic DNNs (DeepSTARR^34^ and BPNet^5^; see Supplementary Table 1). In each benchmark analysis, we compared SQUID to the attribution methods (ISM, Saliency Maps, DeepLIFT, or DeepSHAP) used in the original study describing the DNN being modeled. We found that the attribution maps provided by SQUID consistently yielded binding motifs with markedly-lower attribution variation than the binding motifs provided by the other attribution methods (Fig. 3a). These results were robust to the amount of flanking DNA used when computing attribution variation (Fig. 3b). We also found that strong GE nonlinearities were pervasive in the surrogate models inferred by SQUID for the genomic sequences tested (Extended Data Fig. 3). These results suggest that SQUID, quite generally, quantifies TF binding motifs more consistently than do competing attribution methods. These findings also suggest that modeling GE nonlinearities and heteroscedastic noise is important for the accurate surrogate modeling of genomic DNNs.

**Table 1.** Attribution method performance on a zero-shot variant-effect prediction task. Performance of four attribution methods, applied to three DNNs, on the CAGI5 variant-effect prediction challenge.^47^ Numbers indicate the average correlation observed across the 15 genomic loci assayed by MPRA in ref.^47^. Bold text indicates the best-performing method for each DNN. P-values report results from a paired Wilcoxon signed-rank test on the 15 locus-specific correlation values: *, 0.01 *≤ p* < 0.05; **, 0.001 *≤ p* < 0.01; ***, *p* < 0.001. MPRA, massively parallel reporter assay; DNN, deep neural network; ISM, *in silico* mutagenesis; GE, global epistasis.

| DNN |  | Average Pearson correlation |  |  |  | Statistical significance |  |  |
| --- | --- | --- | --- | --- | --- | --- | --- | --- |
| Architecture | Task | Saliency | ISM | SQUID (Ridge) | SQUID (GE) | GE vs. Saliency | GE vs. ISM | GE vs. Ridge |
| Enformer | DNASE | 0.2977 | 0.4498 | 0.4486 | **0.4801** | *** | ** | ** |
| Basenji-32 | ATAC-seq | 0.2727 | 0.3575 | 0.3700 | **0.4036** | *** | *** | ** |
| ResBind-32 | ATAC-seq | 0.2846 | 0.3388 | 0.3567 | **0.3912** | ** | *** | * |

**Figure 3.**
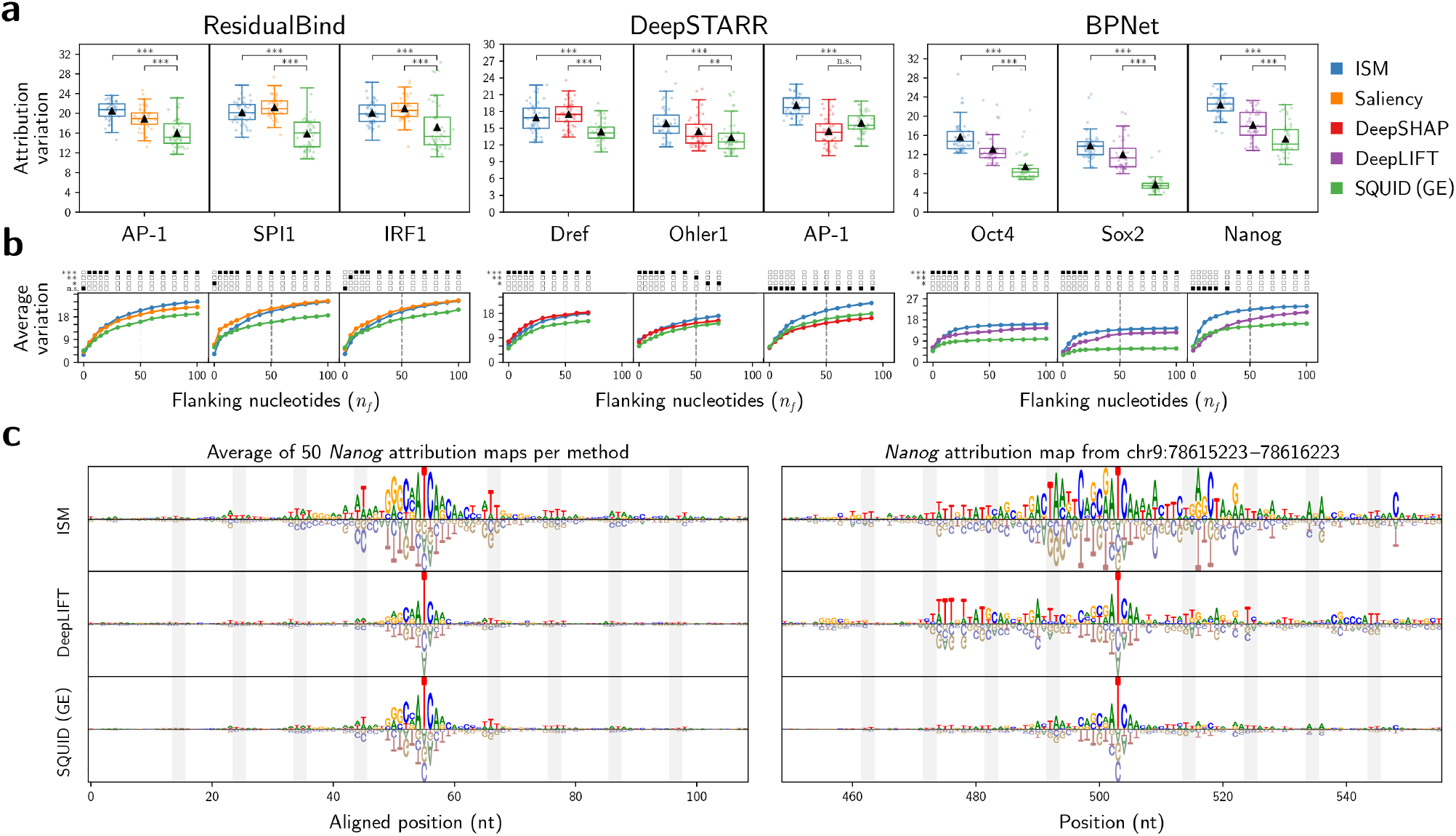
Benchmark analysis across TFs, DNNs, and flank sizes. **a**, Attribution variation analysis for various TFs and genomic DNNs. Results are visualized as in Figure 2b. Each test used 50 sequences from either the human genome (for ResidualBind-32), the mouse genome (for BPNet), or the fly genome (for DeepSTARR), together with consensus TF binding site listed in Supplementary Table 1, and flanked by 50 nt of DNA. **b**, Mean attribution scores computed as in panel **a**, but using variable lengths of flanking DNA. The boxes above each plot indicate the largest (i.e., least significant) p-value, computed as in Figure 2b. n.s., *p ≥* 0.05; *, 0.01 *≤ p* < 0.05; **, 0.001 ≤ *p* < 0.01; ***, *p*< 0.001. Dashed line, flank size used in panel **a. c**, Binding motifs computed for Nanog via attribution analysis of BPNet. *Left*, Attribution maps averaged across 50 mouse loci containing (and centered on) the consensus Nanog binding site AGCCATCAA. *Right*, Attribution map observed for a single such locus. Gray bars indicate 10.5 nt periodicity on either side of the consensus Nanog binding site. TF, transcription factor. DNN, deep neural network. ISM, *in silico mutagenesis*. GE, global epistasis.

The ability of SQUID to better quantify TF binding motifs is exemplified in Figure 3c. Shown are attribution maps for the mouse TF Nanog, computed using BPNet, a consensus binding site of AGCCATCAA, and 50 nt of flanking DNA. When attribution maps are averaged across genomic loci, ISM, DeepLIFT, and SQUID all produce attribution maps that reveal both the Nanog binding motif and a more subtle preference for periodically-spaced AT-rich sequences in the flanking DNA (which likely reflects the interaction of nucleosomes with the DNA double helix). However, the attribution maps for specific genomic sequences often exhibit many other noticeable features. Some of these features are likely to be biological, as TFs often bind in clusters^36^. But many features appear to be spurious and likely reflect non-biological noise in the attribution map. The attribution map provided by SQUID appears to exhibit less noise in the sequences that directly flank core motifs than the attribution maps given by ISM or DeepLIFT. Similar observations hold in analyses of other TFs (Extended Data Fig. 4). These findings raise the possibility that the increased consistency of binding motifs identified by SQUID is due, at least in part, to the reduction of non-biological noise in attribution maps.

### SQUID reduces noise in attribution maps

Non-biological noise in attribution maps can arise from the inherent roughness in the DNN function, a phenomenon that is often associated with benign overfitting.^37, 38^ Benign overfitting is of particular concern for Saliency Maps, since it can adversely affect the accuracy of DNN gradients.^39–41^ Benign overfitting is similarly expected to impact other attribution methods, since these methods essentially quantify how small changes in sequence space affect DNN predictions.^10, 40^ We reasoned that, because SQUID integrates information over an extended region of sequence space, the attribution maps provided by SQUID might be less noisy than the maps provided by other attribution methods.

We therefore asked whether SQUID can reduce the attribution map variation caused by benign overfitting. To answer this question, we trained a DNN to classify ChIP-seq peaks for the human TF GABPA (see Methods) and saved DNN parameters both before benign overfitting (i.e., using early stopping) and after benign overfitting (Fig. 4a). Benign overfitting is apparent from the plateau in validation-set performance that occurs when near-perfect classification performance is achieved on the training set.^42^ We then selected 100 random sequences from the test set, and for each sequence and each of four attribution methods (SQUID, DeepSHAP, SmoothGrad, and Saliency Maps), quantified the differences between the attribution maps obtained using the pre-overfitting DNN parameters versus the post-overfitting DNN parameters. Figure 4b shows that SQUID provided attribution maps that changed substantially less over the course of benign overfitting. Figure 4c illustrates this behavior for a representative sequence. These results support the hypothesis that the attribution maps provided by SQUID are more robust to the adverse effects of benign overfitting than are other attribution methods. These results also suggest that SQUID, more generally, yields attribution maps that have lower noise than maps computed by other attribution methods.

**Figure 4.**
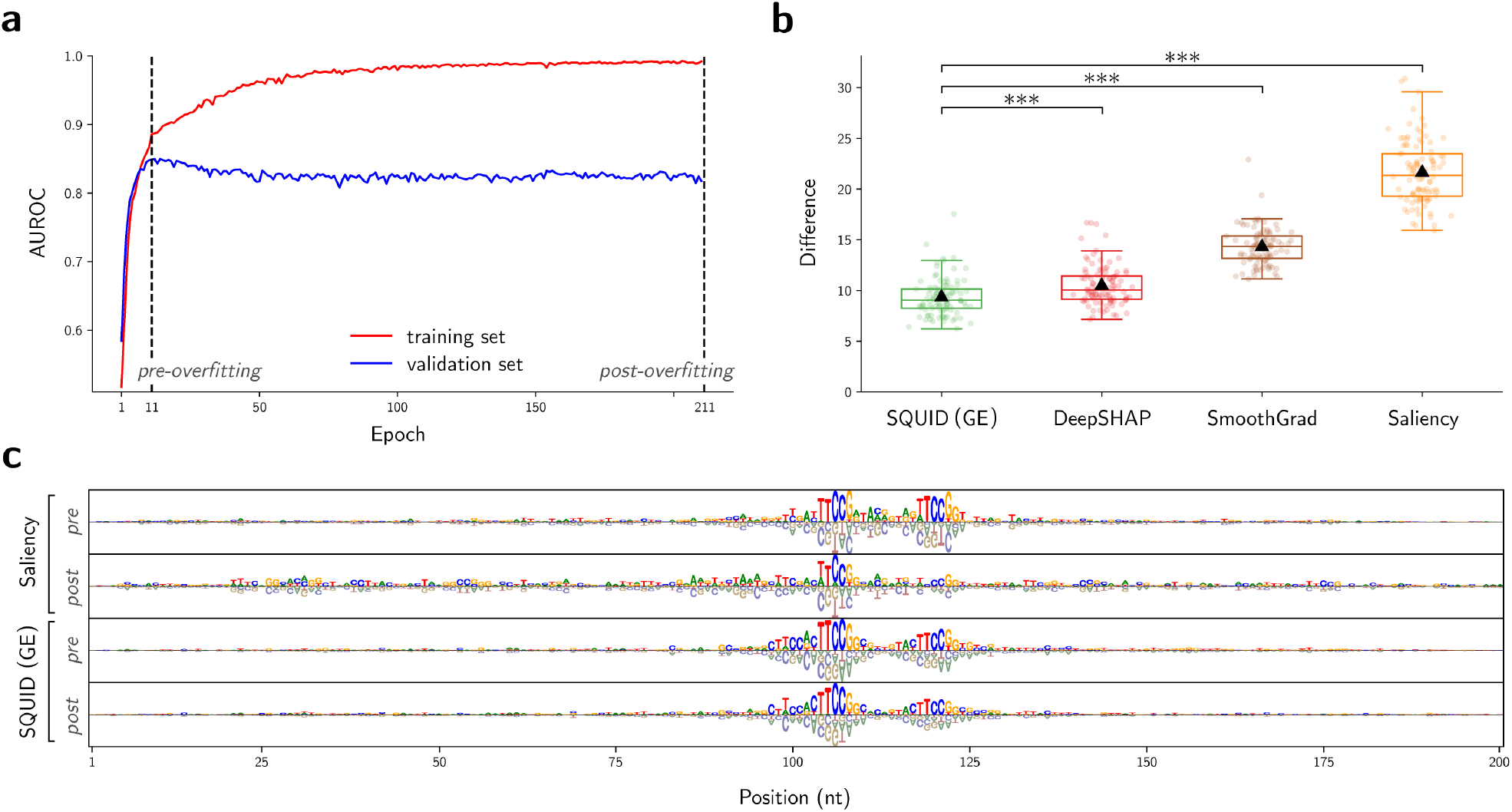
Attribution method performance during benign overfitting. **a**, DNN performance as a function of training epoch. DNN was a 3-layer convolutional neural network trained to classify 200-nt ChIP-seq peaks for the human TF GABPA. Tests used DNN parameters from epoch 11 (pre-overfitting) and epoch 211 (post-overfitting). AUROC, area under the reciever-operator characteristic curve. **b**, Differences between attribution maps for the DNN with pre-overfitting parameters versus post-overfitting parameters, as quantified by the Euclidean distance between attribution map vectors. Results are shown for 100 genomic sequences in the ChIP-seq peak test set. Data is visualized and statistical tests were performed as in Figure 2b. **c**, Attribution maps obtained for a representative test sequence using pre-overfitting and post-overfitting DNN parameters. DNN, deep neural network; TF, transcription factor; GE, global epistasis.

### SQUID better identifies putative weak TF binding sites

Weak TF binding sites play critical roles in eukaryotic gene regulation.^43–45^ Functional signals from weak binding sites, however, can be difficult to distinguish from noise in attribution maps. Having shown that SQUID reduces noise in attribution maps relative to other attribution methods, we hypothesized that SQUID would also better identify weak yet functional TF binding sites. To test this hypothesis, we quantified how well attribution maps generated for putative weak TF binding sites matched the TF binding motifs identified in Figure 3a. For each TF of interest, we randomly selected 150 putative binding sites in the genome having 0, 1, or 2 mutations relative to the consensus binding site (see Supplementary Table 1). We also recorded the genomic sequence containing each selected site padded by 50 nt of flanking DNA on either side. We then computed the score assigned to each selected site using a TF-specific position weight matrix (PWM),^46^ as in Extended Data Figure 5a. Different attribution methods were used to compute attribution maps for each genomic sequence, after which each sequence was assigned an attribution variation value quantified by the Euclidean distance between the attribution map and the corresponding ensemble-averaged attribution map from Figure 3a. Extended Data Figure 5b shows that, as expected, the resulting variation for all attribution methods increased as PWM score decreased. However, the attribution variation observed for SQUID was consistently lower than the attribution variation observed for competing methods. This finding is confirmed by visually examining attribution maps for selected sites (Extended Data Fig. 6). We conclude that SQUID is better than competing methods at identifying signatures of TF binding at weak yet functional TF binding sites.

### SQUID improves the zero-shot prediction of single-nucleotide variant effects

A major goal of genomic DNNs and their attribution methods is to predict which genetic variants are pathogenic. Having observed that SQUID reduces noise in attribution maps, mitigates the adverse effects of benign overfitting, and better identifies TF binding motifs, we hypothesized that SQUID would provide improved variant effect predictions. To test this hypothesis, we used SQUID and other attribution methods to predict the effects that single-nucleotide variants (SNVs) have on the activity of *cis*-regulatory elements for 15 different disease-associated loci in the human genome, loci for which the effects of SNVs had been measured using massively parallel reporter assays (MPRAs).^47, 48^ In technical terms, this is a zero-shot prediction task, since none of the methods tested were trained on MPRA data. At each locus, the Pearson correlation coefficient was computed between the attribution scores and measured SNV effects. We performed this benchmark analysis using three genomic DNNs previously reported for predicting regions of open chromatin: Enformer,^2^ Basenji-32,^49^ and ResidualBind-32.^35^ We found that SQUID yielded substantially higher correlations between predicted and measured SNV effects across the 15 assayed loci (Table 1 and Supplementary Fig. 1). The results suggest that attribution maps provided by SQUID are generally able to predict variant effects better than the attribution maps provided by competing methods. The improved performance of SQUID relative to ISM, which directly uses DNN predictions to quantify SNV effect sizes, suggests that surrogate modeling may provide a general way of improving the variant effect predictions of genomic DNNs themselves.

### SQUID illuminates epistatic interactions

Understanding the role of epistatic interactions within gene regulatory sequences is a major goal in the study of *cis*-regulatory codes. An important advantage of surrogate modeling over other DNN interpretability approaches is that surrogate models with different mathematical forms can be used to answer different questions about DNN behavior. To test whether SQUID could be used to study epistatic interactions in *cis*-regulatory sequences, we implemented a surrogate model that describes all possible pairwise interactions between nucleotides within a sequence (in addition to the additive contributions from individual nucleotides). We then used this pairwise-interaction model to quantify the effects of pairs of putative AP-1 binding sites learned by ResidualBind-32.^35^ We identified 50 genomic sequences having pairs of putative AP-1 binding sites spaced 4 to 20 nt apart, used SQUID to infer the parameters of the pairwise-interaction model about each of the 50 sequences, and then averaged the values of the pairwise-interaction parameters corresponding to both inter-site and intra-site interactions across the 50 sequences (Fig.5a).

The results are shown in Figs. 5b and 5c. We found that pairwise-interaction models consistently performed better on test data than additive surrogate models inferred in a similar manner (Fig.5b). This was true in both the presence and absence of a GE nonlinearity (both additive and pairwise-interaction models benefited from having a GE nonlinearity). The pairwise-interaction models thus provide more accurate approximations of the DNN, suggesting that the pairwise-interaction parameters in these models are likely to be meaningful. Examining the resulting context-averaged pairwise-interaction parameters (Fig. 5c), we observed strong positive intra-site interactions and strong negative inter-site interactions for critical mutations, i.e., mutations away from the preferred nucleotide at any of the 6 most selective positions of the AP-1 motif. Intuitively, the positive interactions between a pair of critical mutations within the same AP-1 site arise because a single critical mutation is sufficient to abrogate AP-1 binding. Thus, a second critical mutation within the same site has no effect, rather than the additional negative effect that would be predicted by an additive model. By contrast, the negative inter-site interactions between critical mutations indicate a degree of redundancy between nearby AP-1 binding sites, since mutations that abrogate both AP-1 sites result in lower activity than is expected from adding the effects of abrogating each AP-1 site individually.^50^

**Figure 5.**
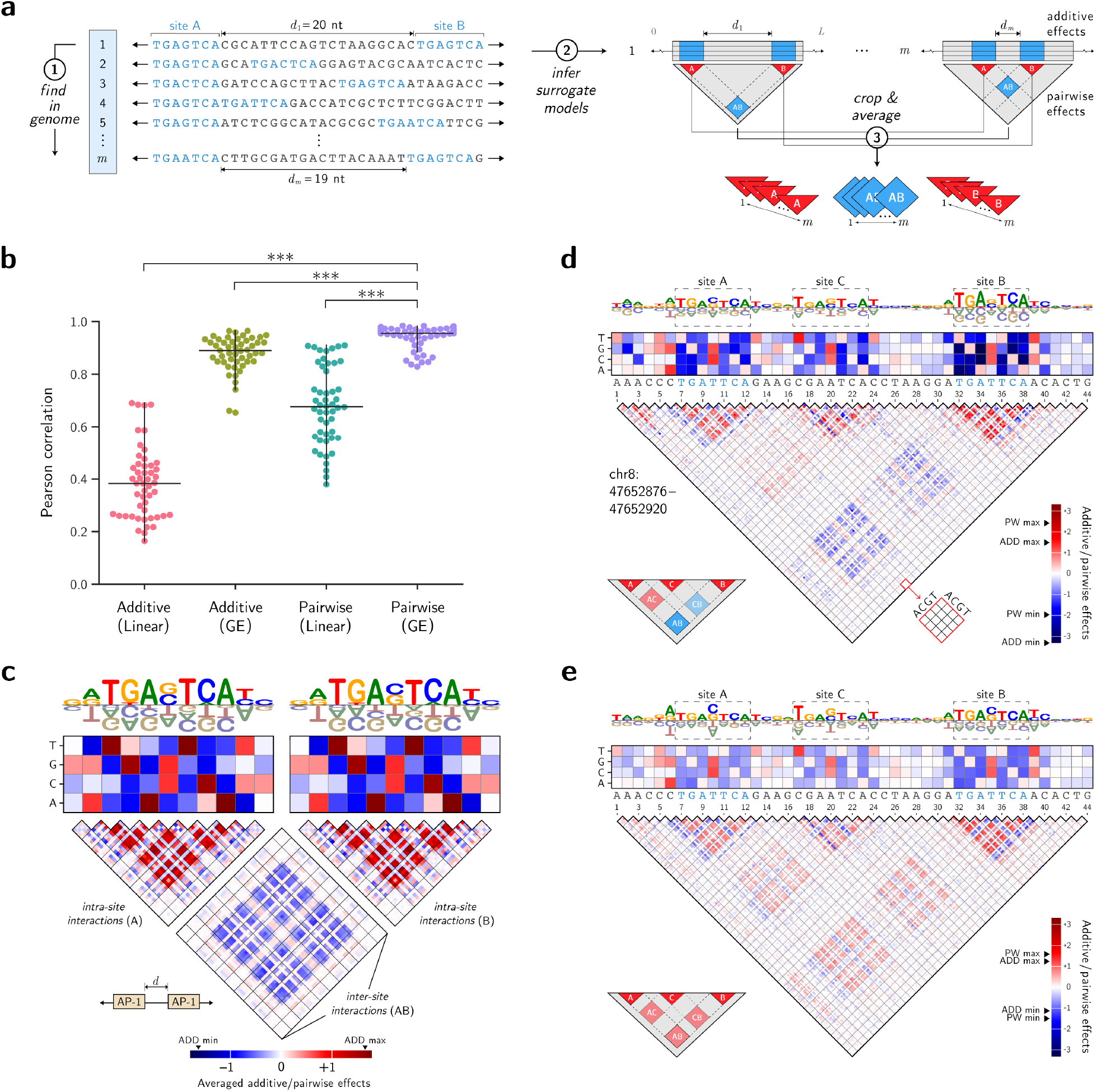
SQUID captures epistatic interactions. **a**, Pipeline for analyzing context-averaged epistatic interactions. The pipeline consists of 3 steps: (1) pairs of consensus binding sites (A and B) are identified in genomic sequences; (2) pairwise-interaction models are inferred for each identified sequence; and (3) the values of surrogate model parameters describing intra-site interactions (A, B) and inter-site interactions (AB) are averaged across sequence contexts. **b**, Performance of surrogate models for 50 genomic sequences having two consensus AP-1 binding sites each. Results are shown for both additive and pairwise-interaction models, each with (GE) and without (linear) a GE nonlinearity. Correlation values were computed between surrogate model predictions and DNN predictions on *in silico* test data. Overlaid lines represent the median, upper, and lower quartiles. P-values were computed using the one-sided Mann-Whitney U test. ***, *p* < 0.001. **c**, Surrogate model parameters quantifying intra-site and inter-site interactions, averaged across the 50 genomic sequence contexts. Pairwise-interaction models having GE nonlinearities were used in this analysis. **d**,**e**, Parameters of pairwise-interaction models, either with (panel **d**) or without (panel **e**) a GE nonlinearity, determined for a genomic locus having three putative AP-1 binding sites. GE, global epistasis; DNN, deep neural network.

While these averages are informative about overall trends, analyses of specific sequences can provide additional insight. In the analysis above, we observed one sequence that contained 3 putative AP-1 binding sites, with different combinations of these sites displaying either positive or negative epistatic interactions (Fig.5d). This example shows that context-specific interactions can be complex, and that other factors like binding site orientation and spacing may play an important role.^51^ We note that including the GE nonlinearity in the pairwise-interaction model was essential for identifying these complex interactions. When a linear pairwise-interaction model was used, we instead observed positive interactions of similar magnitude between every pair of sites (Fig.5e). The reason is that, in the linear pairwise model, the pairwise-interaction parameters are co-opted to describe a global nonlinearity instead of the nucleotide-specific interactions they are intended to model. Specifically, for this genomic sequence, the DNN exhibits a rapidly saturating effect of removing AP-1 binding sites on the functional score (Extended Data Fig. 7), such that disrupting a second binding site has a much smaller effect on DNN predictions than would be predicted based on the effect of disrupting just one of the three sites. We thus see that modeling GE nonlinearities is essential when interpreting genomic DNNs in terms of epistatic interactions between genetic elements.

### SQUID supports global DNN interpretations

In the above analyses, SQUID used *in silico* MAVE libraries generated by partially mutagenizing a specific sequence of interest in the genome. As a result, the surrogate models inferred by SQUID provided DNN interpretations that are limited to localized regions of sequence space. In previous work, however, we proposed a DNN interpretation method called global importance analysis (GIA),^52^ in which the DNN is evaluated on completely random sequences containing embedded genetic elements, such as putative TF binding sites. We therefore investigated whether SQUID could provide useful global DNN interpretations using a modified version of the sequence libraries used in GIA.

First we asked whether SQUID could provide global interpretations for individual TFs. For each TF of interest, we generated an *in silico* MAVE library by embedding partially mutagenized versions of the consensus TF binding site within random DNA (Extended Data Fig. 8a). We then used this *in silico* MAVE library to infer an additive surrogate model. This approach was applied to the four mouse TFs (Oct4, Sox2, Klf4, and Nanog) modeled by BPNet.^5^ Extended Data Figure 8b shows that the resulting additive surrogate models revealed prominent motifs having low background, which matched the known sequence preferences of the four TFs. In the case of Nanog, this analysis also revealed periodic secondary motif features in flanking DNA nearly identical to those in Figure 3c. We conclude that global surrogate modeling with SQUID can provide accurate characterizations of TF binding motifs independent of specific genomic sequence contexts.

We next investigated whether SQUID could provide global insights into the epistatic interactions between pairs of TF binding sites. We created an *in silico* MAVE library in which partially-mutagenized version of the Nanog and Sox2 consensus binding sites were inserted into random DNA sequences a fixed distance apart (Extended Data Fig. 8c). We then used SQUID to infer a pairwise-interaction model based on this *in silico* MAVE library. Extended Data Figure 8d shows that, similar to the findings in Figure5, critical mutations within Nanog and Sox2 binding sites exhibited positive intra-site epistatic interactions and negative inter-site epistatic interactions. We then repeated this analysis for Nanog and Sox2 binding sites separated 0 to 32 nt and observed that the strength of inter-site epistatic interactions varied in a sinusoidal manner consistent with the periodicity of the DNA double helix (Extended Data Fig. 8e). We conclude that global surrogate modeling with SQUID can provide useful characterizations of epistatic interactions between TFs in a manner that is independent of genomic sequence context.

## Discussion

Here we introduced SQUID, a framework for interpreting genomic DNNs. SQUID uses surrogate models to approximate DNN functions in user-defined regions of sequence space. The parameters of these surrogate models can then be mechanistically interpreted: additive surrogate models can be interpreted as attribution maps, and pairwise-interaction surrogate models can be interpreted as quantifying epistatic interactions. Applying SQUID to a variety of genomic DNNs, we observed that the attribution maps obtained by SQUID more robustly identify TF binding motifs and provide better variant-effect predictions than the attribution maps obtained using other DNN interpretability methods. We also observed that SQUID is able to quantify epistatic interactions that are otherwise obscured by global nonlinearities in the DNN.

SQUID works by using the DNN of interest as a forward simulator of MAVE experiments. SQUID then uses the MAVE-NN modeling framework^26^ to infer surrogate models from the resulting *in silico* data. This surrogate modeling approach has multiple important advantages over standard DNN interpretability methods.

First, SQUID is model agnostic: it does not require access to the parameters or gradients of the DNN. Rather, SQUID simply uses the DNN of interest as a black-box oracle. This allows SQUID to be applied to arbitrary genomic DNNs, regardless of their computational implementation.

Second, SQUID smooths over fluctuations in DNN predictions that are likely to reflect noise rather than biological function. This is because the parameters of the surrogate models that SQUID infers are fit to DNN predictions over an extended region of sequence space. Our results showed that this increased smoothness can cause the surrogate models to have improved accuracy relative to the parent DNN.

Third, SQUID leverages domain-specific knowledge to improve the utility of surrogate models. While the use of surrogate models for interpreting DNNs has been established previously in other fields, these applications typically use linear models.^28^ By using the latent phenotype models supported by MAVE-NN, the surrogate models inferred by SQUID explicitly account for global nonlinearities and heteroscedastic noise, both of which are ubiquitous in functional genomic data and in genomic DNNs. By explicitly accounting for these influences, SQUID is able to remove their confounding effects on the inferred surrogate model parameters.

Finally, SQUID supports a variety of surrogate models. In this work we demonstrated the use of additive and pairwise-interaction models, but SQUID also supports use of surrogate models having user-specified mathematical form. SQUID thus has the ability to infer surrogate models that reflect higher-order epistatic interactions,^21, 22, 25, 53^ or specific biophysical hypotheses (e.g. thermodynamic models^54–58^ or kinetic models^59–62^). We note, however, that more research is still needed into how to best distill biological signals from confounding noise in attribution maps.

Although nonlinearities and genetic interactions can be identified by observing whether changes to sequence cause changes to additive attribution maps (e.g., maps provided by Saliency Maps, DeepLIFT, or DeepSHAP), explicitly accounting for these nonlinearities and genetic interactions within the surrogate model allows SQUID to quantify these effects in a natural way. One limitation to modeling epsitatic interactions in SQUID using pairwise-interaction surrogate models is that the number of such interactions, and thus the number of parameters that must be inferred, increases quadratically with sequence length. We found that, in practice, the inference of pairwise-interaction models presented computational difficulties when input sequences were longer than approximately 100 nt. The number of parameters in additive models increases only linearly with sequence length, and such models are often more practical to implement on long sequences as a result.

The biggest drawback of SQUID relative to other DNN interpretability methods (other than LIME, which has the same drawback) is its higher computational demands. This is due to the need for many forward passes through the DNN during simulation of the *in silico* MAVE dataset, as well as the need to fit the surrogate model parameters to these simulated data (see Extended Data Table 1 for typical SQUID computation times). We therefore suggest that SQUID is likely to be more useful for in-depth analyses of specific sequences of interest (e.g., disease-associated loci) rather than large-scale genome-wide analyses. In this context, however, the advantages of SQUID described above–especially SQUID’s ability to support surrogate models of arbitrary mathematical form–enables new ways of biologically interpreting genomic DNNs.

## Methods

### The SQUID framework

An overview of SQUID’s workflow is given in Supplementary Figure 2. Briefly, SQUID takes as input a sequence of interest and a specified surrogate model. An *in silico* MAVE dataset is generated with the `InSilicoMAVE` object, with specifications of the mutagenesis strategy given by a `Mutagenizer` object, and processed model predictions given by a `Predictor` object. The *in silico* MAVE dataset is then fit with surrogate models which are defined as objects in `squid/surrogate_zoo.py`.

- **Mutagenizer**. An *in silico* MAVE dataset is generated by sampling a library of sequences using random partial mutagenesis of a sequence of interest. We modulate the size of the sequence-space region from which this library is drawn using two hyperparameters: the sequence that defines the region of interest, which has length *L*, and the mutation rate *r*. The resulting number of mutations in each individual sequence is a Poisson distributed random variable having mean *Lr*. `squid/mutagenizer.py` contains objects that apply the chosen mutagenesis strategy, with the object `RandomMutagenesis` executing the random partial mutagenesis in this study.
- **Predictor**. SQUID currently requires that DNNs provide scalar-valued outputs. However, some genomic DNNs output high-dimensional predicted profiles, not scalar predictions. For example, ResidualBind-32 predicts 15 chromatin accessibility profiles of length 64, each profile corresponding to a different cell type and each profile element representing a binned position with a resolution of 32 nt. `squid/predictor.py` contains objects, including `ScalarPredictor` and `ProfilePredictor`, that reduce model predictions to scalar values. For profile-based predictions, SQUID also offers an approach to reduce profiles using principal component analysis (PCA). Specifically, profiles are projected onto their first principal component, with sign chosen so that the wild-type sequence has higher-than average score (see Supplementary Fig. 3).
- **InSilicoMAVE**. The object `InSilicoMAVE`, defined in `squid/mave.py`, is a data structure that takes the mutagenizer and predictor objects and generates the *in silico* MAVE dataset for a sequence of interest.
- **Surrogate Zoo**. The `squid/surrogate_zoo.py` module currently offers a linear model (`SurrogateLinear`) as well as models based on MAVE-NN (`SurrogateMAVENN`). In `SurrogateLIME`, the “K-Lasso” approach is implemented.^28^ First, `sklearn` regression is performed using `linear_model.LassoLars` to model the entire regularization path over all values of *α*. We then select the model associated with the appropriate *α* that gives the desired number of nonzero features, *K*; i.e., weights. These *K* features are then fit using least squares linear regression. In `SurrogateRidgeCV`, a linear model is computed using `sklearn` ridge regression with iterative fitting along a regularization path, where the best model is selected by cross-validation (`linear_model.RidgeCV`). `SurrogateMAVENN` supports models based on the built-in modeling capabilities of MAVE-NN. The mathematical forms of these models are described in detail in ref.^26^

### Surrogate models

The surrogate models supported by MAVE-NN for use in SQUID comprise three parts (Fig. 1b): a G-P map, a GE nonlinearity, and a noise model. Here we summarize the mathematical forms of these model components. In what follows, **x** represents a sequence of interest, *x*_*l*:*c*_ is a one-hot encoding of **x** (i.e., is equal to 1 if **x** has character *c* at position *l* and is equal to 0 otherwise), and *y* is the DNN-predicted scalar activity of **x**.

- The **G-P map**, *f* (**x**, ***θ***), which has parameters ***θ*** , maps a sequence **x** to a latent phenotype *ϕ* . There are two types of G-P maps: additive G-P maps and pairwise-interaction G-P maps. Additive G-P maps have a constant parameter *θ*_0_, additive parameters *θ*_*l*:*c*_, and the mathematical form
- 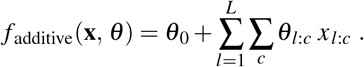
- Pairwise-interaction G-P maps have a constant parameter, additive parameters, pairwise-interaction parameters *θ*_*l*:*c*,*l*_^*′*^_:*c*_^*′*^ , and the mathematical form
- 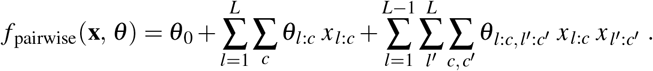
- The **GE nonlinearity**, *g*(*ϕ*), maps the latent phenotype *ϕ* to a most-probable scalar DNN prediction *ŷ*. By default, *g*(*ϕ*) is defined to be an over-parameterized linear combination of hyperbolic tangent sigmoids. In this paper we use 50 sigmoids, corresponding to 50 hidden nodes in the neural network formulation of *g*(*ϕ*). This is the default number of sigmoids used by MAVE-NN, and was chosen based on previous analyses of diverse MAVE datasets^26^. We found that TF motifs inferred by SQUID, as well as corresponding *R*^2^ values, were highly robust to the number of hidden nodes (Extended Data Table 2). For SQUID analyses performed without a GE nonlinearity, *g*(*ϕ*) is defined to be a linear function of *ϕ* .
- The **noise model**, *p*(*y*| *ŷ*), describes the expected distribution of DNN predictions *y* about the most-probable prediction *ŷ*. The noise model can be defined using a Gaussian distribution, a Student’s t-distribution, or the skewed t-distributed of Jones and Faddy.^30^ The shape parameters of this distribution can either be independent of *ŷ* (for homoscedastic noise) or a polynomial function of *ŷ* (for heteroscedastic noise).

### LIME

LIME is a surrogate modeling approach for interpreting general DNNs. Here we describe the general LIME framework, how SQUID generalizes LIME, and how the two implementations of LIME used for the benchmarking results in Figure 2 and Extended Data Figure 1 work.

Let *x* denote an one-hot encoded sequence input, *y*(*x*) denote the prediction of the DNN of interest, *π*(*x*) denote a probability distribution on sequence space, *f* (*x, θ*) denote the surrogate model used to approximate *y*, and *θ* denote the parameters of the surrogate model. We note that ref.^28^ distinguishes between the input *x* fed to the DNN and the “interpretable input representation” *x*^*′*^ used as input to the surrogate model inferred by LIME. In our genomics applications, however, we take both *x* and *x*^*′*^ to be the standard one-hot encoding of sequence. We therefore omit this distinction in what follows.

LIME determines values for surrogate model parameters via

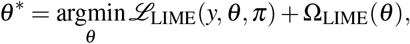

where *ℒ*_LIME_ is a loss function and Ω_LIME_ is a regularization penalty on *θ* . Following ref.^28^, we adopt the weighted *L*_2_ loss function

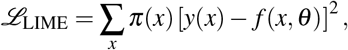

which is estimated by randomly sampling inputs *x* according to the probability distribution *π*(*x*).

SQUID can be thought of as a generalization of LIME in which the loss function has latent parameters that must be inferred simultaneously with the parameters of the surrogate model. As in LIME, the G-P map *f* (*x, θ*) serves as the surrogate model. But unlike LIME, the SQUID loss function is determined by a measurement process *p*_MP_(*y*| *f* , *η*), which represents a probability distribution over DNN predictions *y* conditioned on the prediction *f* of the G-P map, and which has its own set of latent parameters (denoted by *η*). SQUID simultaneously determines values for *θ* and *η* via

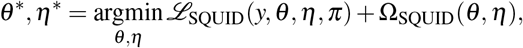

where Ω_SQUID_ is a regularization penalty on both *θ* and *η*, and the loss function *L*_SQUID_ is given by

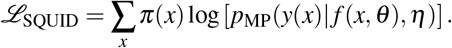

In terms of the GE nonlinearity *ŷ* ( *f* , *η*) and the noise model *p*_noise_(*y*| *ŷ, η*), the measurement process is given by

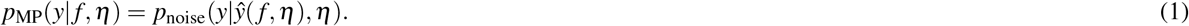

In Main Text we consider two different implementations of LIME: Ridge and K-Lasso. The Ridge implementation is equivalent to SQUID when the GE nonlinearity is taken to be the identity function and the noise model is taken to be a Gaussian with fixed variance. In this implementation, we also take Ω_LIME_ to be an *L*_2_ penalty with scaling coefficient determined by cross-validation. The K-Lasso implementation is equivalent to the specific algorithm describe in ref.^28^, if *π*(*x*) is taken to be the distribution in sequence space resulting from a Poisson mutation process. In this implementation, the user selects a value *K* for the number of nonzero one-hot features (as DNA sequence of length *L* having 4*L* such features). Lasso regression is then used to select the *K* specific features for which *θ* is allowed to be nonzero, and the specific values of these nonzero elements of *θ* are inferred by least squares.

### Deep learning models

This study used six DNNs: ResidualBind-32,^35^ Basenji-32,^35^ DeepSTARR,^34^ Enformer,^2^ BPNet,^5^ and a baseline CNN that predicts ChIP-seq data for the human TF GABPA. Here we briefly describe each DNN and how that DNN was used in our study to compute a prediction *y* for each sequence **x** when generating *in silico* MAVE data.

- **ResidualBind-32** predicts ATAC-seq profiles across 15 human cell lines.^35^ ResidualBind-32 takes as input a DNA sequence of length 2048 nt and outputs 15 profiles (one for each cell line) where each profile comprises 64 bins, with each bin spanning 32 nt. The published ResidualBind-32 parameters were used to compute these profiles. In our attribution variation analyses, *y* was set equal to the sum of predicted binned ATAC-seq signals over all 64 bins for the single output channel corresponding to the cell type most associated with the TF of interest (see Supplementary Table 1). In our variant-effect analysis, *y* was set equal to the sum of predicted binned ATAC-seq signals over all 64 bins, using a profile averaged across all 15 output channels.
- **Basenji-32** predicts ATAC-seq profiles across 15 human cell lines.^49^ The input and output of Basenji-32 is identical to that of ResidualBind-32, and the published Basenji-32 parameters were used to computed predicted ATAC-seq profiles. Analyses performed using Basenji-32 were performed the same way as for ResidualBind-32.
- **DeepSTARR** predicts *Drosophila* enhancer activity as assayed by UMI-STARR-seq.^34^ DeepSTARR takes as input a DNA sequence of length 249 nt and outputs two scalar-valued predictions for enhancer activity for developmental (Dev) and housekeeping (Hk) regulatory programs. The published DeepSTARR parameters were used to predict enhancer activity. In each analysis, *y* was computed using the regulatory program most associated with the TF of interest (see Supplementary Table 1).
- **Enformer** predicts many different types of functional genomic tracks (e.g., ChIP-seq, DNase-seq, ATAC-seq, and CAGE) across the human and mouse genomes.^2^ Enformer takes as input a DNA sequence of length 393,216 nt and (for humans) outputs 5,313 profiles (one for each track) where each profile comprises 128 bins, each bin spanning 32 nt, representing the central 114,688 nt of the input sequence. The published Enformer parameters were used to compute these profiles. In our variant-effect analysis, we used human predictions, where *y* was computed as in the original study by cropping all 674 “cell-type agnostic” DNase profiles to a 10-bin (1280 nt) region centered about the variant of interest, then summing across bins in the mean cropped profile.
- **BPNet** predicts nucleotide-resolution ChIP-nexus binding profiles for four TFs (Oct4, Sox2, Klf4, and Nanog) in mouse embryonic stem cells.^5^ BPNet takes as input a DNA sequence of length 1000 nt and outputs a 1000-valued positive (+) and negative (-) strand profile for each of the four TFs (8 profiles in total per input). The parameters of BPNet were retrained as specified in the original release,^63^ and the resulting model was confirmed to recapitulate the published model `BPNet-OSKN`^64^ by a visual inspection of attribution maps. Analyses for different prediction tasks used different profiles (see Supplementary Table 1). In all BPNet analyses except those in Extended Data Figure 8, *y* was computed using the profile contribution score defined in the original paper.^5^ In the BPNet analysis for Extended Data Figure 8, *y* was instead computed using PCA as described above and in Supplementary Figure 3.
- The **baseline CNN** predicts ChIP-seq peaks for the human TF GABPA in GM12878 cells. This model takes as input a DNA sequence of length 200 nt and outputs a single probability. In our analysis of the effects of benign overfitting, *y* was computed as the logit of the output probability.

The baseline CNN has not been previously published. Briefly, this model was trained to distinguish binary-labeled sequences: ChIP-Seq peaks of GABPA in human GM12878 cells (positive labels) and DNase-seq peaks of GM12878 cells that did not overlap with any GABPA peaks (negative labels). Data was acquired from https://zenodo.org/record/7011631 (data/GABPA_200.h5); 11,022, 1,574, and 3,150 sequences in the training, validation, and test sets were respectively used. The CNN takes as input a DNA sequence of length 200 nt and outputs a probability. The hidden layers consist of three convolutional blocks each with max pooling (size 4 with stride 4), followed by a fully-connected hidden layer with 128 units and an output layer to a single node with sigmoid activations. The number of filters and the kernel sizes from the first convolutional layer to the third are: (32, 15), (64, 5), and (96, 5). The same padding was used for each convolutional layer. All hidden layers used ReLU activations and dropout,^65^ with rates given in order by 0.1, 0.2, 0.3, and 0.5. The CNN was trained to minimize the binary cross-entropy loss function using Adam^66^ with default parameters and a batch size of 64. Early stopping was implemented to save model parameters at the epoch that corresponded to the largest area-under the ROC curve (AUROC) on the validation set, yielding the pre-overfitting DNN parameters. From that point, the CNN was trained for an additional 200 epochs, yielding the post-overfitting DNN parameters.

### Attribution methods

Our analyses used attribution maps computed using a variety of methods, implemented as follows.

- ***In silico* mutagenesis (ISM) scores** were computed by evaluating the scalar DNN prediction for every single-nucleotide variant of the sequence of interest.
- **Saliency Maps scores** were computed by evaluating the gradient of the scalar DNN prediction at the sequence of interest with respect to the one-hot encoding of that sequence.
- **DeepSHAP scores** were computed as previously described for DeepSTARR.^34^
- **DeepLIFT scores** were computed as previously described for BPNet.^5^
- **SmoothGrad scores** were computed by averaging Saliency Maps over 50 noisy encodings of the sequence of interest. Each noisy encoding was computed by adding Gaussian noise (mean zero, standard deviation 0.25) to each of the 4*L* matrix elements of the one-hot encoding for the sequence of interest.

### Standardization of attribution maps

Prior to comparing attribution maps, we standardized these maps to remove non-identifiable and/or non-meaningful degrees of freedom. Using *v*_*l*:*c*_ to denote the attribution map value for character *c* at position *l*, this standardization was carried out as follows.

- For **plotting sequence logos** and **computing attribution variation values**: Attribution map values were standardized using the transformation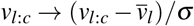, where 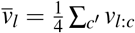 and 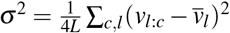. This transformation is essentially a gradient correction at each position,^67^ followed by a normalization with the square root of the total variation of the attribution scores across the sequence.
- For **variant-effect analysis** (Table 1): Attribution map values were standardized using the transformation 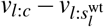, where *s*^wt^ is the wild-type sequence and 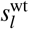 is the character at position *l* in this sequence.
- For **plotting heatmaps** of additive and pairwise-interaction model parameters (Fig. 5c,d,e and Extended Data Fig. 8d): Parameters were standardized as in ref.^26^ using the “empirical gauge”.

### Attribution variation computations

Attribution variation values were computed as follows.

- For **consensus TF binding sites** (Fig. 2 and Fig. 3), we first located all instances of the consensus TF binding site in the genome; the consensus sites used for each TF are listed in Supplementary Table 1. Using a baseline attribution method (Saliency Maps for ResidualBind-32, ISM for DeepSTARR and BPNet), we then ranked and manually pruned these genomic sequences to identify *m* = 50 putative functional and spatially-isolated TF binding sites. For each of the *m* genomic sequences, we then computed an attribution map spanning the putative TF binding site plus 100 nucleotides on either side (i.e., flanks). We then cropped these *m* attribution maps to span *n* _*f*_ nucleotides on either side of the consensus TF binding site. For Figure 2 we used *n* _*f*_ = 15; for Figure 3 we used *n* _*f*_ = 0, 5, 10, 20, 30,…, 100. For each of the *m* cropped attribution maps, the attribution variation was defined to be the Euclidean distance between the cropped attribution map and the average of the *m* cropped attribution maps.
- For **weak TF binding sites** (Extended Data Fig. 5), we first located all instances of variants of the consensus TF binding site in the genome having up to 2 naturally-occurring mutations. For each group of putative binding sites having 0, 1, or 2 mutations, we then ranked and manually pruned these genomic sequences to identify *m* = 50 putative functional and spatially-isolated TF binding sites as above. For each of the 3*m* genomic sequences, we then computed an attribution map spanning the putative TF binding site plus 100 nucleotides on either side. We then cropped these 3*m* attribution maps to span *n* _*f*_ = 50 nucleotides on either side of the consensus TF binding site. For each of the 3*m* cropped attribution maps, the attribution variation was defined to be the Euclidean distance between the cropped attribution map and the average of the *m* cropped attribution maps from the ensemble of binding sites having 0 mutations.

### Context-averaged epistatic interactions

To compute the context-averaged epistatic interactions shown in Figure 5c, first, we located all pairs of consensus AP-1 binding sites in the genome separated by no more than 20 nucleotides. We then ranked and pruned these pairs to identify *m* = 50 genomic sequences for further analysis. For each of the *m* genomic sequences, we used SQUID to infer a pairwise-interaction model spanning the pair of putative sites and 6 nucleotides on either side. Parameter values were standardized by expressing them in the empirical gauge.^26^ Additive parameters for each site were then cropped and averaged over the *m* sequences. The intra-site and inter-site pairwise-interaction parameters were similarly cropped and averaged. This process is schematized in Figure 5a.

### Variant-effect analysis

The ability of different attribution methods to predict variant effects, quantified in Table 1, was benchmarked as follows.

- The **variant-effect data** used to perform these benchmark studies is MPRA data from ref.^47^. These data comprise measurements for the effects of nearly all SNVs in 15 disease-associated regulatory sequences, with each sequence ranging from 187 nt to 600 nt in length. For each assayed SNV, variant effect was quantified as the difference in measured activity between the variant and wild-type sequences.

To compute **attribution map predictions of variant effects**, we extracted the genomic sequences centered about each of the 15 assayed regulatory sequences (2,048 nt per sequence for tests of ResidualBind-32 and Basenji-32; 393,216 nt per sequence for tests of Enformer). For each attribution method, the effect of an SNV, from character 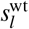 to character *c* at position *l* in each regulatory sequence *s*^wt^, was quantified as 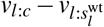. When inferring surrogate models using SQUID, we mutagenized the regulatory sequence and 400 nt of flanking sequence on either side.

## Data availability

The datasets, models, and computational results used to support the findings in this paper are available on Zenodo at https://doi.org/10.5281/zenodo.10047748.^68^ Datasets include the test set sequences held out during the training of ResidualBind-32, DeepSTARR, and BPNet; the ChIP-seq peaks and background sequences used to train our 3-layer CNN; and the CAGI5 challenge dataset.

## Code availability

SQUID is an open-source Python package based on TensorFlow.^69^ SQUID can be installed via pip (https://pypi.org/project/squid-nn) or GitHub (https://github.com/evanseitz/squid-nn). Documentation for SQUID is provided on ReadTheDocs (https://squid-nn.readthedocs.io). The code for performing all analyses in this paper is available on GitHub as well (https://github.com/evanseitz/squid-manuscript), and a static snapshot of this code is available on Zenodo.^68^

## Acknowledgements

We thank Amber Tang, Shushan Toneyan, Mahdi Kooshkbaghi, Chandana Rajesh, Jakub Kaczmarzyk, and Carlos Martí for helpful discussions. This work was supported in part by: the Simons Center for Quantitative Biology at Cold Spring Harbor Laboratory; NIH grants R01HG012131 (PKK, ESS, JBK, DMM), R01HG011787 (JBK, ESS, DMM), R01GM149921 (PKK), R35GM133777 (JBK), and R35GM133613 (DMM); and an Alfred P. Sloan Foundation Research Fellowship (DMM). Computations were performed using equipment supported by NIH grant S10OD028632.

## Author contributions

ESS, DMM, JBK, and PKK conceived of the study. EES wrote the software and performed the analysis. ESS designed the analysis with help from DMM, JBK, and PKK. JBK and PKK supervised the study. ESS, DMM, JBK, and PKK wrote the manuscript.

---

**Extended Data Table 1.**
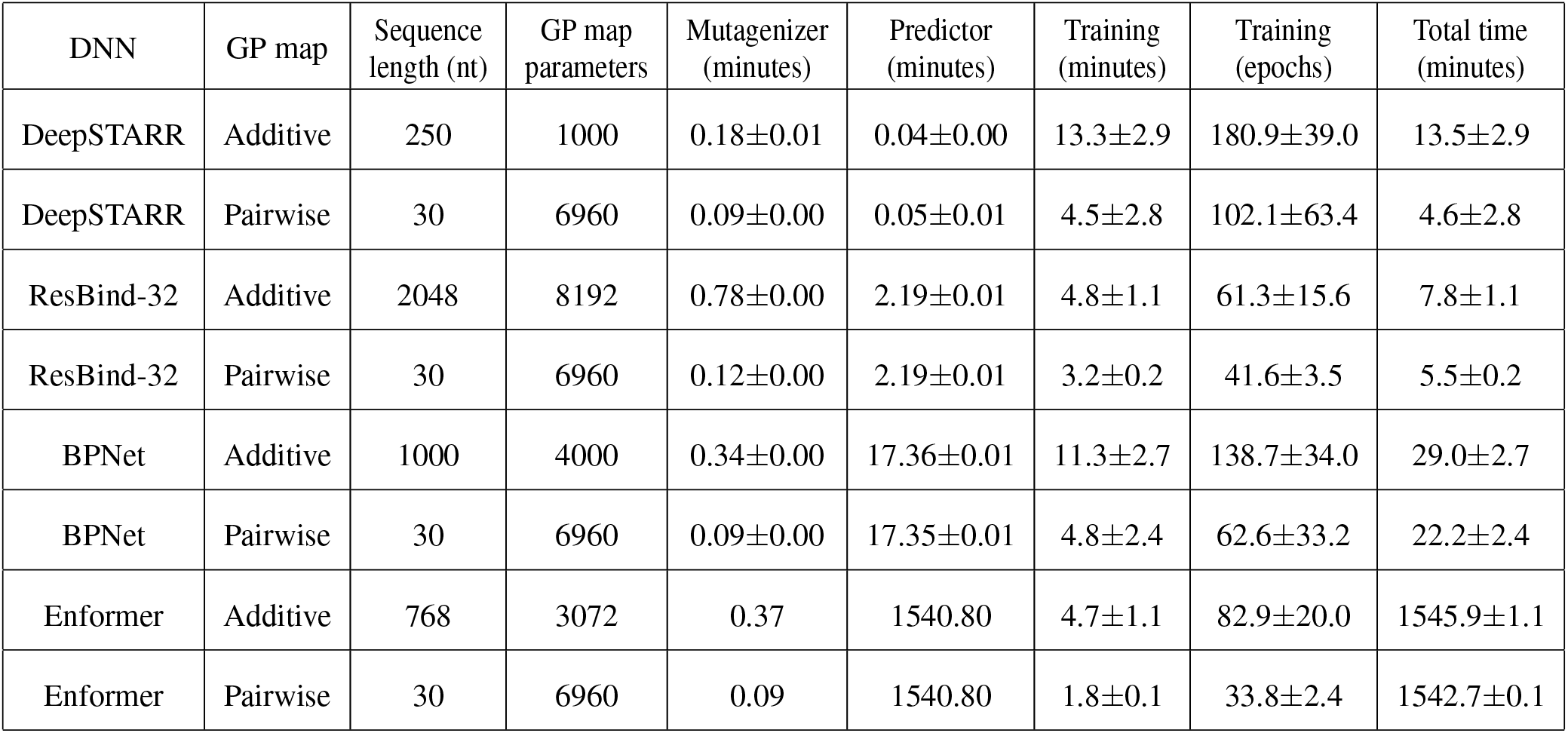
SQUID computation times. Shown are SQUID computation times for the different stages of analysis of various genomic DNNs using various types of surrogate models. All analyses were performed using 100K variant sequences mutagenized at 10% per nucleotide, a GE measurement process with a skewed-T noise model, and early stopping. Error bars indicate standard deviations over 10 analysis runs. For DeepSTARR, ResBind-32, and BPNet, analyses were performed for 10 different in silico MAVE datasets. For Enformer, 10 analyses were performed on the same in silico MAVE dataset due to each forward-pass being slower for Enformer than for the other DNNs. Shorter sequences were used when inferring pairwise-interaction models to keep the number of pairwise-interaction model parameters comparable to the number of additive model parameters. DeepSTAR and ResBind-32 analyses were performed using a V100 GPU in Google Colab. BPNet and Enformer analyses were performed using an RTX A4000 GPU on a private server.

| DNN | GP map | Sequence length (nt) | GP map parameters | Mutagenizer (minutes) | Predictor (minutes) | Training (minutes) | Training (epochs) | Total time (minutes) |
| --- | --- | --- | --- | --- | --- | --- | --- | --- |
| DeepSTARR | Additive | 250 | 1000 | 0.18±0.01 | 0.04±0.00 | 13.3±2.9 | 180.9±39.0 | 13.5±2.9 |
| DeepSTARR | Pairwise | 30 | 6960 | 0.09±0.00 | 0.05±0.01 | 4.5±2.8 | 102.1±63.4 | 4.6±2.8 |
| ResBind-32 | Additive | 2048 | 8192 | 0.78±0.00 | 2.19±0.01 | 4.8±1.1 | 61.3±15.6 | 7.8±1.1 |
| ResBind-32 | Pairwise | 30 | 6960 | 0.12±0.00 | 2.19±0.01 | 3.2±0.2 | 41.6±3.5 | 5.5±0.2 |
| BPNet | Additive | 1000 | 4000 | 0.34±0.00 | 17.36±0.01 | 11.3±2.7 | 138.7±34.0 | 29.0±2.7 |
| BPNet | Pairwise | 30 | 6960 | 0.09±0.00 | 17.35±0.01 | 4.8±2.4 | 62.6±33.2 | 22.2±2.4 |
| Enformer | Additive | 768 | 3072 | 0.37 | 1540.80 | 4.7±1.1 | 82.9±20.0 | 1545.9±1.1 |
| Enformer | Pairwise | 30 | 6960 | 0.09 | 1540.80 | 1.8±0.1 | 33.8±2.4 | 1542.7±0.1 |

**Extended Data Table 2.** GE modeling is robust to the number of hidden nodes. Shown are results for SQUID analyses performed using surrogate models having either no GE nonlinearity (Linear) or a GE nonlinearity (GE) computed using different numbers of hidden nodes (i.e., tanh sigmoidal components; see Methods). Analyses were performed as described in Extended Data Table 1, except that in each test the 10 in silico MAVEs were generated for the same genomic locus (as opposed to different genomic loci). For DeepSTARR, an additive GP-map covering 250 nt was inferred. For ResBind-32, an additive GP-map covering 2048 nt was inferred. Error bars represent standard deviations over the 10 in silico MAVE datasets.

| DNN | Measurement Process | GE Hidden Nodes | $R^2$ | Epochs | test I_pred (bits) | max I_var (bits) | max val_I_var (bits) |
| --- | --- | --- | --- | --- | --- | --- | --- |
| DeepSTARR | Linear | N/A | 0.70±0.03 | 98.4±23.4 | 1.2±0.1 | 0.5±0.1 | 0.5±0.1 |
| DeepSTARR | GE | 1 | 0.86±0.03 | 112.4±24.7 | 1.3±0.1 | 1.2±0.1 | 1.3±0.3 |
| DeepSTARR | GE | 50 | 0.86±0.03 | 202.1±41.5 | 1.3±0.1 | 1.3±0.1 | 1.2±0.1 |
| DeepSTARR | GE | 1000 | 0.86±0.03 | 146.8±54.7 | 1.3±0.1 | 1.3±0.1 | 1.2±0.1 |
| ResBind-32 | Linear | N/A | 0.70±0.18 | 113.8±37.8 | 1.4±0.2 | 0.2±0.4 | 0.0±0.4 |
| ResBind-32 | GE | 1 | 0.93±0.04 | 136.9±79.4 | 2.3±0.2 | 2.4±0.2 | 2.0±0.2 |
| ResBind-32 | GE | 50 | 0.94±0.04 | 116.7±80.3 | 2.3±0.2 | 2.4±0.2 | 2.0±0.3 |
| ResBind-32 | GE | 1000 | 0.93±0.05 | 127.0±47.8 | 2.3±0.2 | 2.4±0.2 | 2.1±0.2 |

**Extended Data Figure 1.**
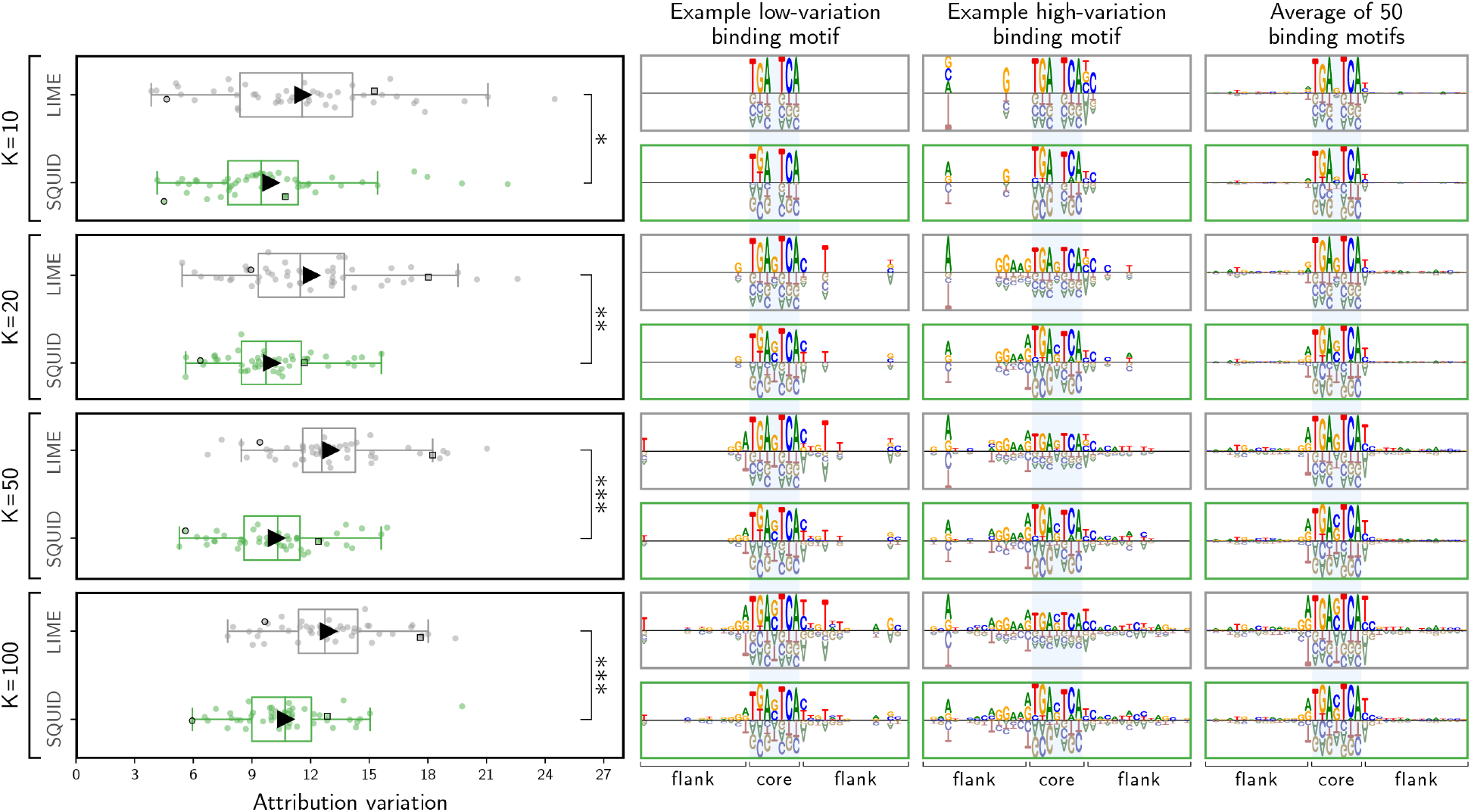
Additional comparisons of K-Lasso LIME to SQUID. Shown are the results of analyses, performed as in Fig. 2b, comparing the performance of SQUID to the performance of the K-Lasso implementation of LIME for four different values of *K*. p-values were computed using a one-sided Mann-Whitney U test; ***, *p* < 0.001. We note that the attribution variation values obtained for SQUID in these tests varied systematically with the choice of *K*. The reason is as follows. The K-Lasso LIME algorithm produces sparse attribution maps that have only *K* nonzero parameters. Consequently, the variation observed in K-Lasso LIME attribution maps systematically decreases as *K* decreases. This gives K-Lasso LIME an unfair advantage in the attribution variation test described in Main Text and in Methods. To fairly compare K-Lasso LIME to SQUID in this figure, we therefore modified this test. In the analysis of each in silico MAVE, the attribution map elements inferred by SQUID were first set to zero at the same positions where all K-Lasso LIME attribution map elements were exactly zero. Attribution variation values were then calculated as described in Main Text and in Methods.

**Extended Data Figure 2.**
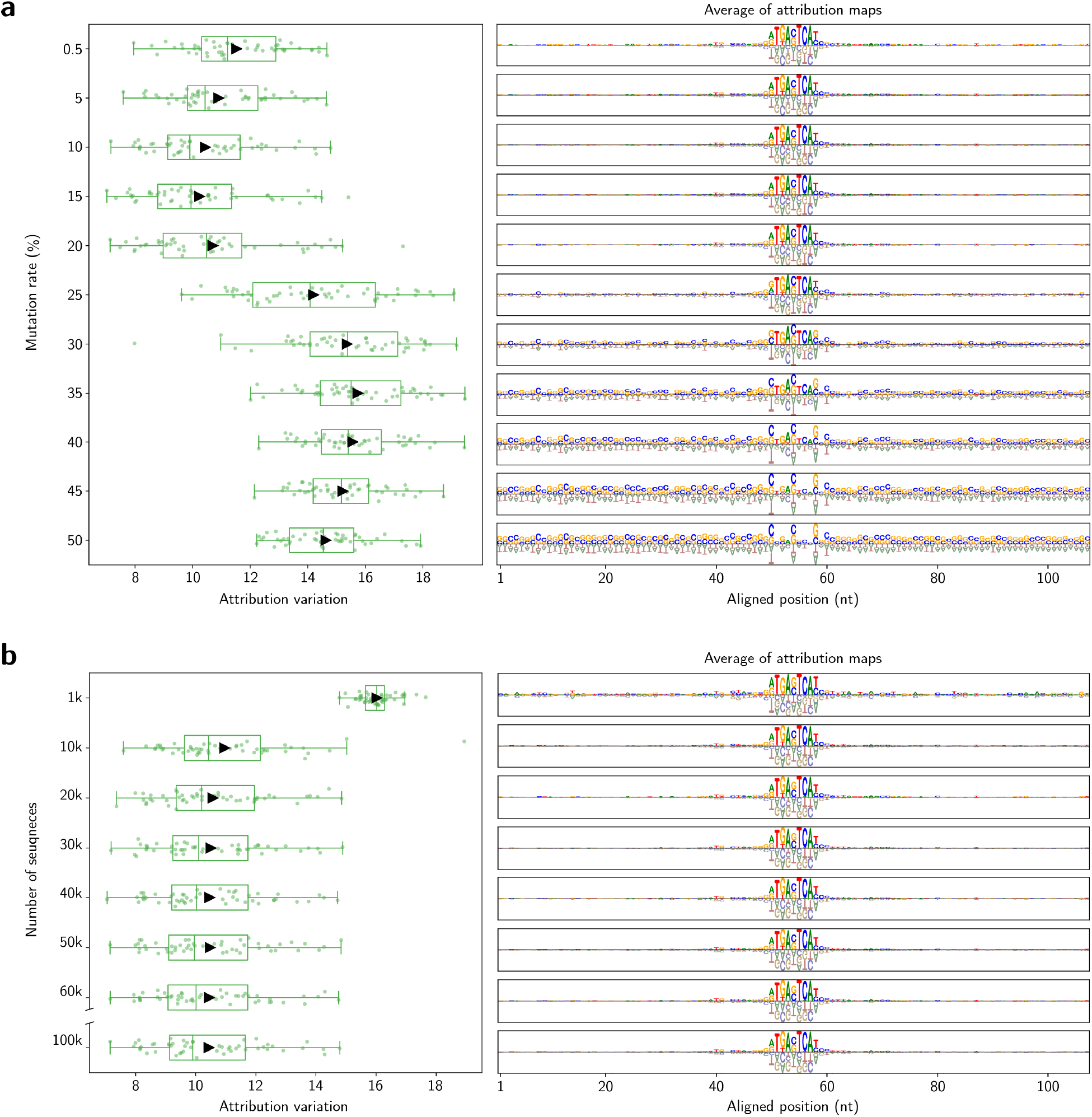
Influence of mutation rate and library size on SQUID attribution maps. **a** and **b**, attribution variation (*left*) and average attribution maps (*right*) found for 50 SQUID attribution maps computed for the TF AP-1 as in Fig. 2, but using *in silico* MAVE libraries having (**a**) variable mutation rate *r* and fixed size *N* = 100, 000, or (**b**) fixed mutation rate *r* = 10% and variable size *N*. All SQUID attribution maps were computed using additive models with GE nonlinearities, followed by cropping these maps using a flank size of *n* _*f*_ = 50 nt. TF, transcription factor; MAVE, multiplex assay of variant effect; GE, global epistasis.

**Extended Data Figure 3.**
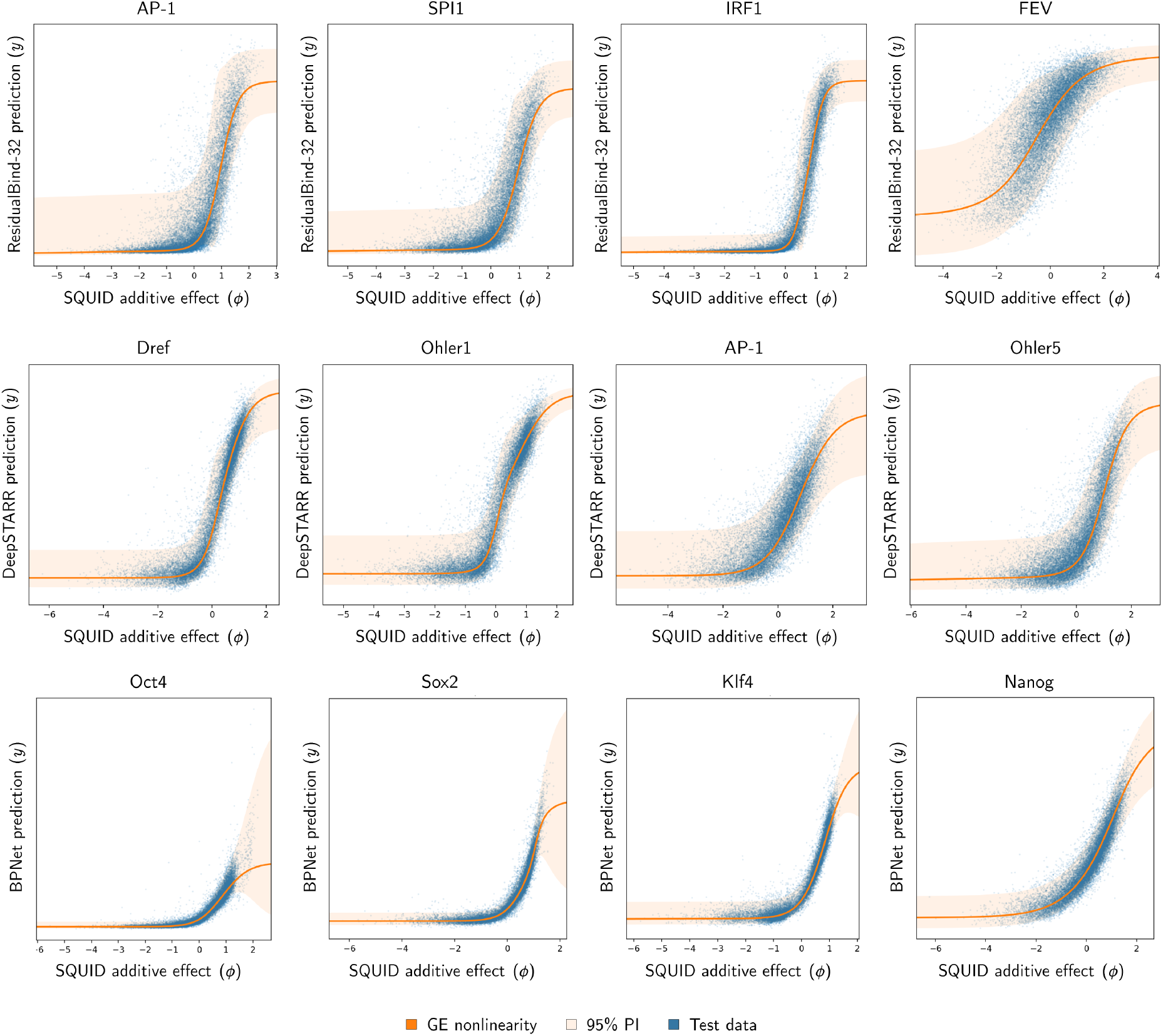
Nonlinearities and noise across DNNs and TFs. Examples of the GE nonlinearities and heteroscedastic noise models inferred in the SQUID (GE) analyses performed for Fig. 3. Each plot shows results for a representative sequence from the 50 sequences analyzed for each combination of DNN and TF. DNN, deep neural network; TF, transcription factor; GE, global epistasis; PI, prediction interval.

**Extended Data Figure 4.**
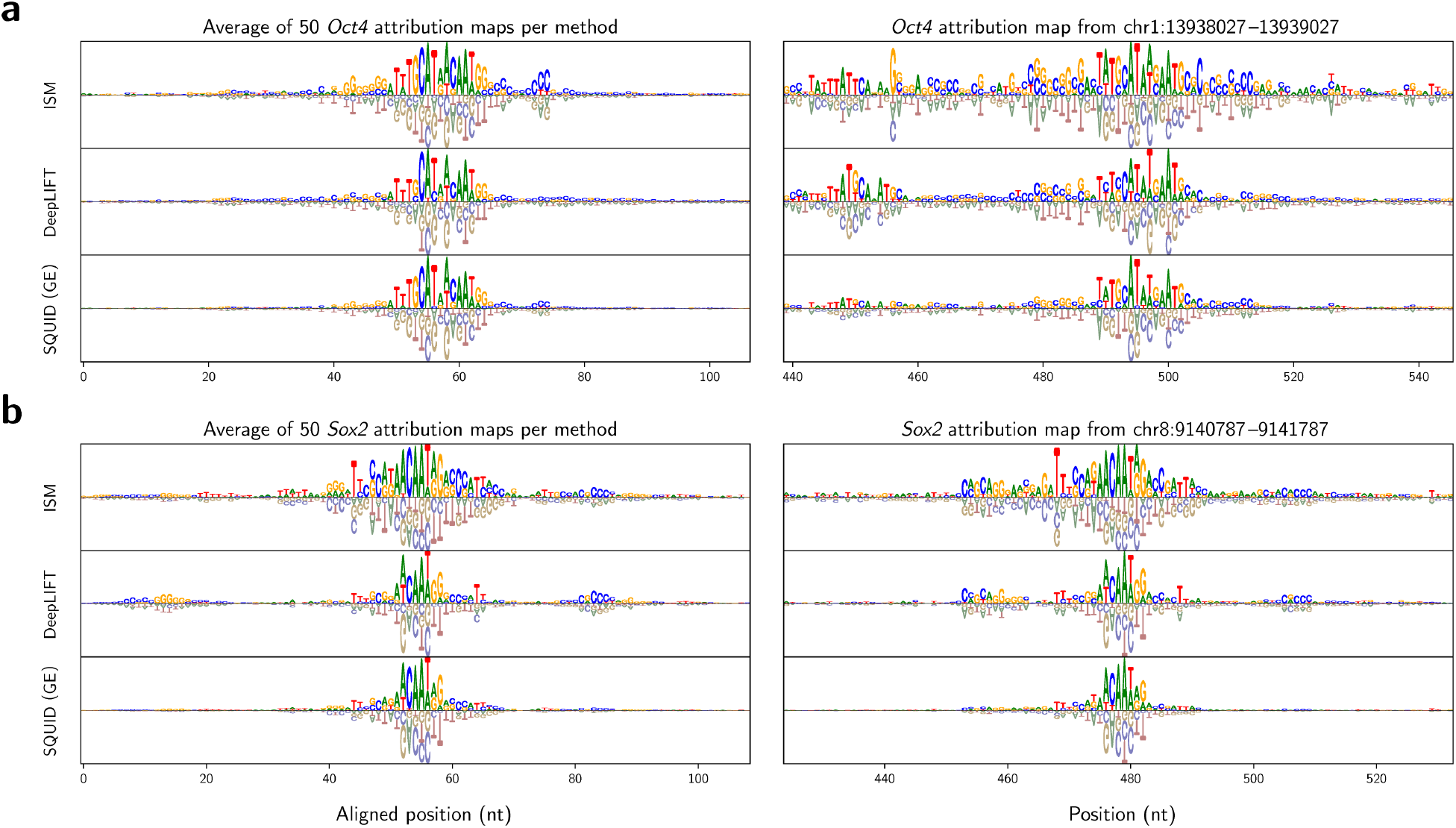
Average and example binding motifs for Oct4 and Sox2. **a**, Oct4 motifs, centered on the putative binding site `TTTGCAT`. **b**, Sox2 motifs, centered on the putative binding site GAACAATAG. TF binding motifs are from attribution maps computed for BPNet and plotted as in Fig. 3c. TF, transcription factor.

**Extended Data Figure 5.**
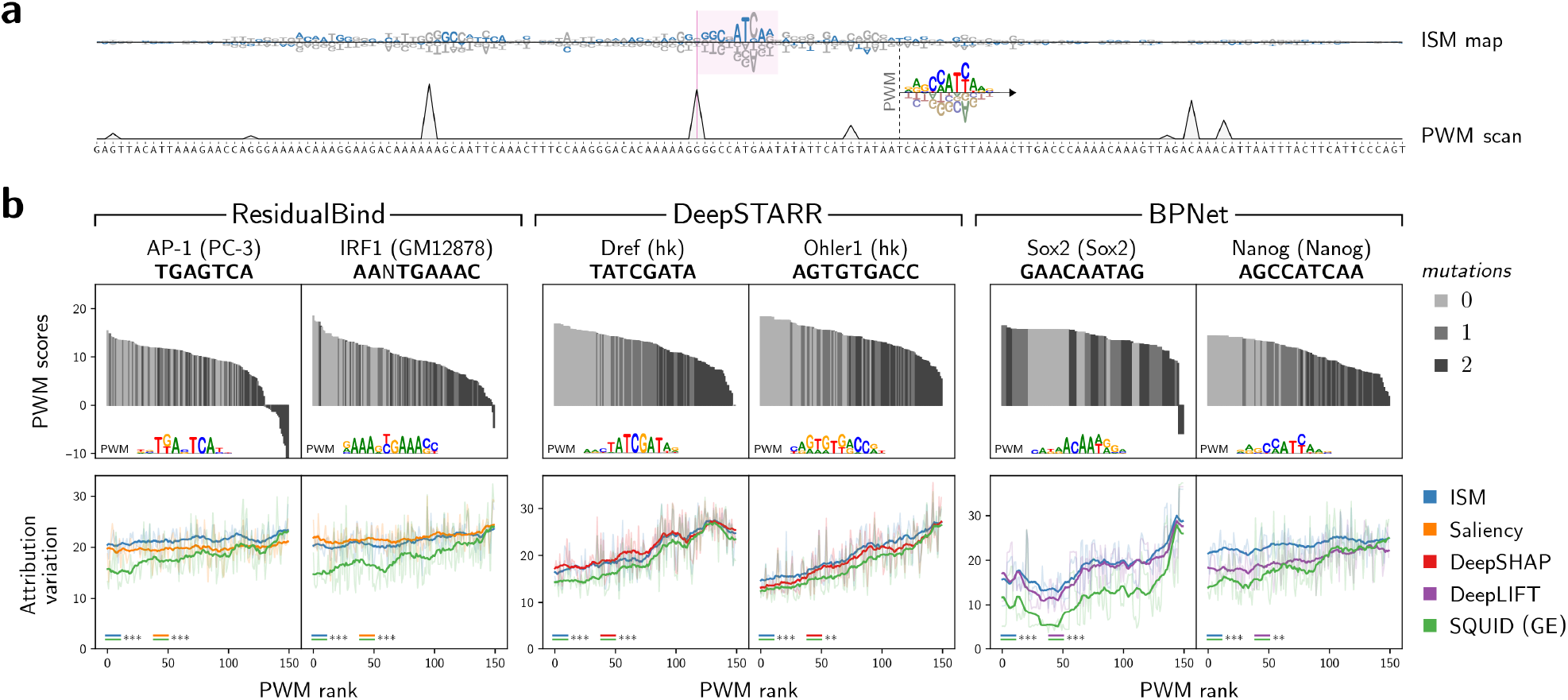
Benchmark analysis of attribution variation for putative weak TF binding sites. **a**, *Upper*, Attribution map for a representative genomic sequence containing multiple putative weak binding sites for the mouse TF Nanog. Attribution map is for the BPNet DNN and was computed using ISM. Blue, wild-type nucleotides; gray, non-wild-type nucleotides. *Middle*, PWM for Nanog. *Lower*, PWM scores across the genomic sequence. Only positive PWM scores are shown. **b**, For each TF and DNN, plots show attribution variation values for 150 putative TF binding sites plotted against putative binding site strength. Bold lines indicate signals smoothed with a sliding window of 20 nt. Stars indicate p-values computed using the one-sided Mann-Whitney U test: **, 0.001 *≤ p* < 0.01; ***, *p* < 0.001. PWM scores for each of the 150 putative sites are shown above, along with a logo representation of the PWM used. Each site is represented by a gray bar shaded according to the number of mutations (0, 1, or 2) in the core of the putative site.

**Extended Data Figure 6.**
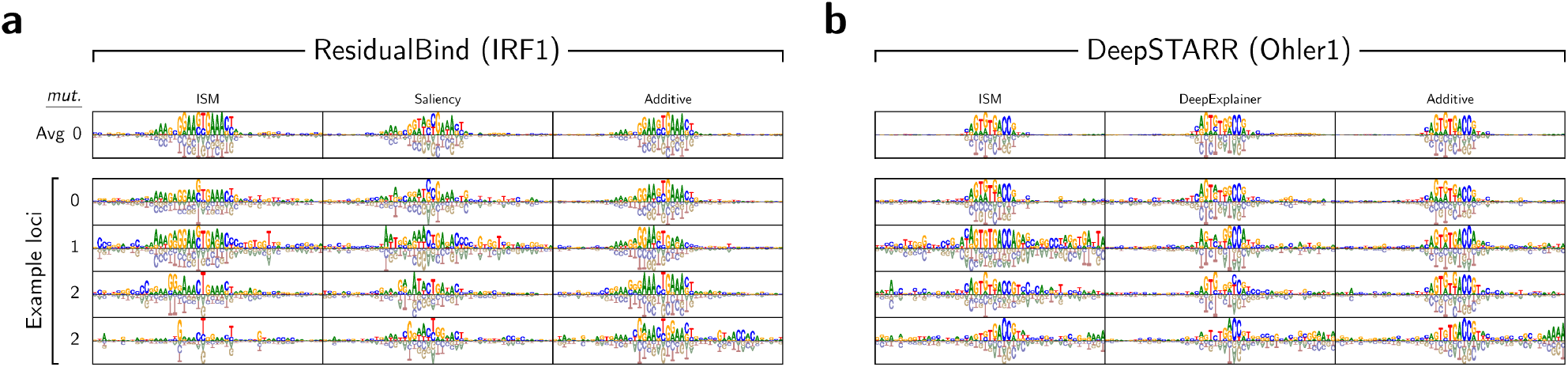
Attribution maps computed for strong and weak TF binding sites. Top row shows the average of 50 attribution maps in the 0-mutation ensemble computed for (**a**) IRF1 using ResidualBind-32, and for (**b**) Ohler using DeepSTARR. Remaining rows show attribution maps for four representative genomic loci with the central putative binding site having varying numbers of mutations from the consensus binding site. TF, transcription factor.

**Extended Data Figure 7.**
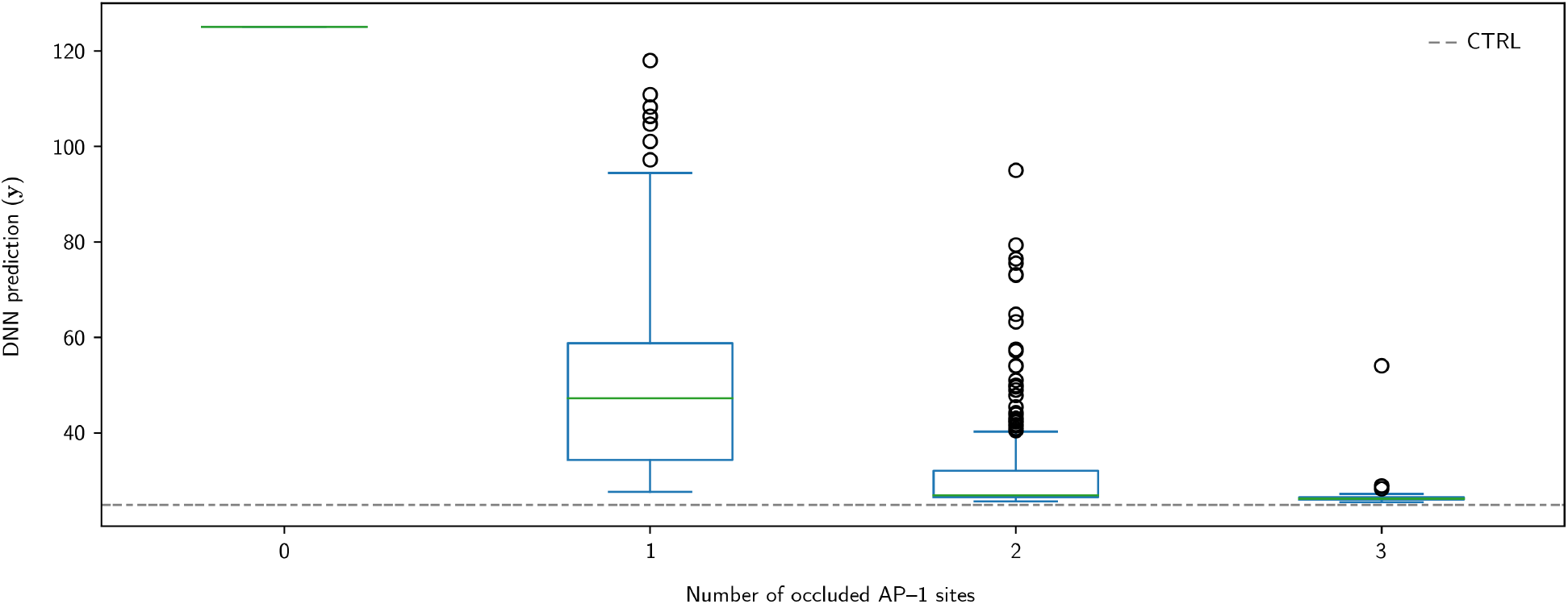
Occlusion analysis of AP-1 binding site effects. Occlusion study based on the wild-type sequence investigated in Fig. 6d and 6e. Occlusions were performed 100 times for every combination of one, two, or three occluded motifs, with the DNN prediction taken independently for each instance. In each occluded sequence, the corresponding AP-1 core (7-mer) sites were scrambled using a uniform probability of nucleotides at each position. The baseline score (CTRL) was calculated from the median of predictions corresponding to 100 instances of a dinucleotide shuffle over the full (2048-nt) sequence. The DNN prediction rapidly approaches the genomic baseline as additional binding sites are occluded.

**Extended Data Figure 8.**
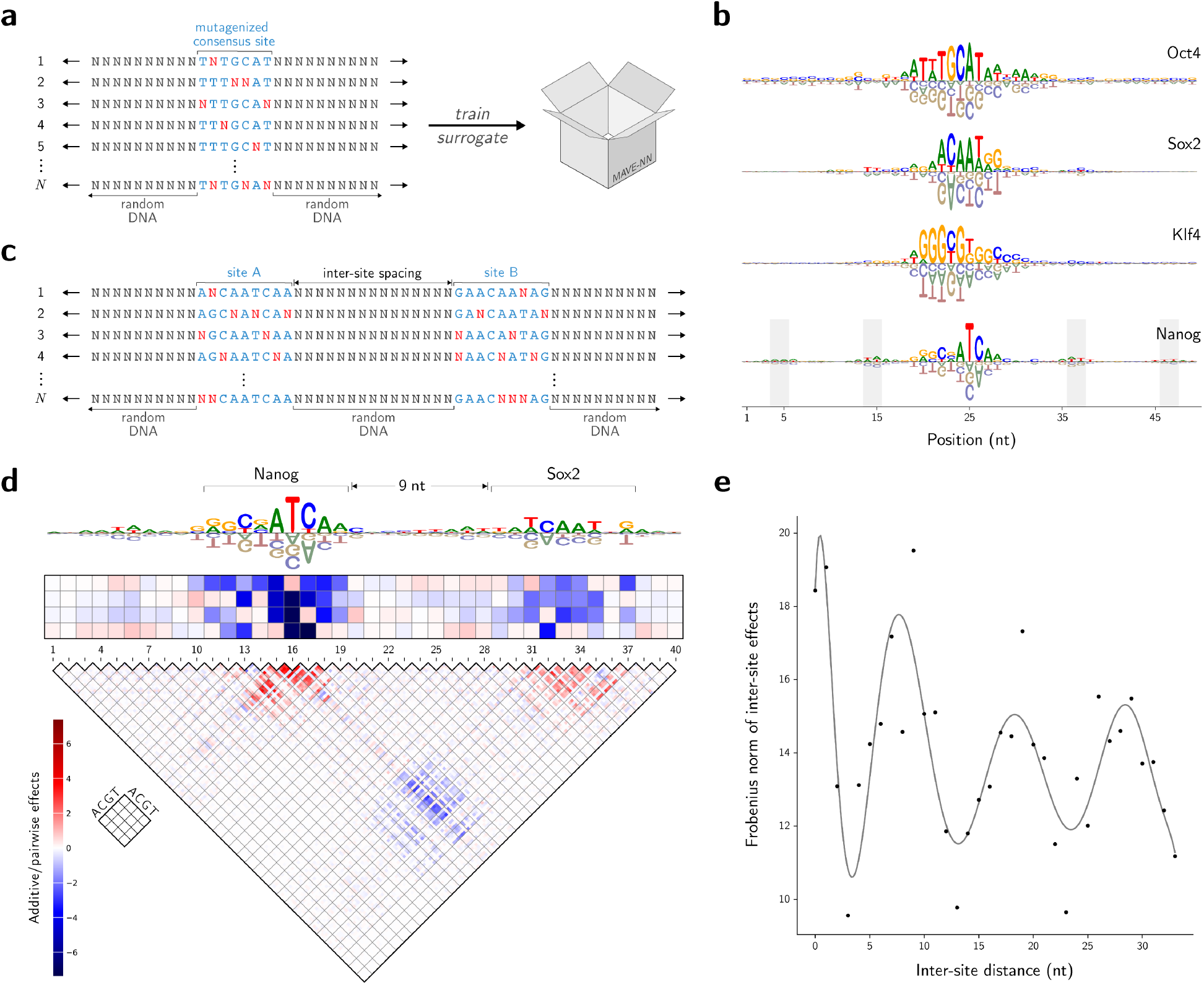
SQUID supports global DNN interpretations. **a**, *In silico* MAVE libraries used by SQUID to infer surrotate models of global TF specificity. Each sequence in the library contains a partially-mutagenized version of a consensus TF binding site embedded within random DNA. **b**, Additive surrogate models for BPNet^?^ inferred by SQUID using libraries of the form in panel **a** for the mouse TFs Oct4, Sox2, Klf4, and Nanog. Gray bars indicate 10.5-nt periodicity on either side of the inferred Nanog motif. **c** Format of the *in silico* MAVE libraries used to study global TF-TF epistatic interactions. Each sequence in the library contains two partially mutagenized consensus TF binding sites embedded a fixed distance apart (0 nt to 32 nt) within random DNA. **d**, Pairwise surrogate model inferred for BPNet by SQUID using a library of the form in panel c and putative binding sites for Nanog and Sox2. **e**, Distance-dependence of inter-site interactions between Nanog and Sox2. Dots show the Frobenius norm of inter-site pairwise interactions, inferred as in panel **d**, using libraries having different distances between the embedded Nanog and Sox2 sites. The occurrence of periodic inter-site interaction minima occurred at distances where the central `AA` in Sox2 overlapped with the periodic Nanog-associated flanking signals. Black line is from the least-squares fitting of a polynomial of degree 10. Libraries used for panels **d** and **e** comprised 500,000 sequences each. BPNet profiles were projected to scalars using PCA, which yielded substantially lower background noise than obtained using profile contribution scores. DNN, deep neural network; MAVE, multiplex assay of variant effect; TF, transcription factor.

**Supplementary Table 1.** TFs analyzed in our study. Shown for each TF is the consensus sequence used, the DNN that models the TF, the DNN prediction task to which attribution methods were applied, and PWMs used to investigate weak binding sites. TF, transcription factor; DNN, deep neural network; PWM, position weight matrix.

| TF name | consensus seq. | DNN name | DNN task | PWM ID (database) |
| --- | --- | --- | --- | --- |
| AP-1 | TGAGTCA | ResBind-32 | PC-3 | MA0476.1 (Jaspar) |
| AP-1 / AP-1 | TGANTCA ... TGANTCA | ResBind-32 | PC-3 | n/a |
| IRF1 | TGAAAC | ResBind-32 | GM12878 | n/a |
| IRF1-long | AANTGAAAC | ResBind-32 | GM12878 | MA0050.1 (Jaspar) |
| SPI1 | GGAAGT | ResBind-32 | GM12878 | n/a |
| FEV | CCGGAA | ResBind-32 | HCT116 | n/a |
| AP-1 | TGACTCA | DeepSTARR | Dev | MA0476.1 (Jaspar) |
| Dref | TATCGATA | DeepSTARR | Hk | M00230 (Homer) |
| Ohler1 | AGTGTGACC | DeepSTARR | Hk | M00232 (Homer) |
| Ohler5 | CAGCTG | DeepSTARR | Hk | n/a |
| Oct4 | TTTGCAT | BPNet | Oct4 | n/a |
| Sox2 | GAACAATAG | BPNet | Sox2 | H12CORE.0.P.B (Hocomoco) |
| Klf4 | GGGTGTGGC | BPNet | Klf4 | n/a |
| Nanog | AGCCATCAA | BPNet | Nanog | H12CORE.1.P.B (Hocomoco) |
| Nanog / Sox2 | AGCCATCAA ... GAACAATAG | BPNet | Nanog | n/a |

**Supplementary Figure 1.**
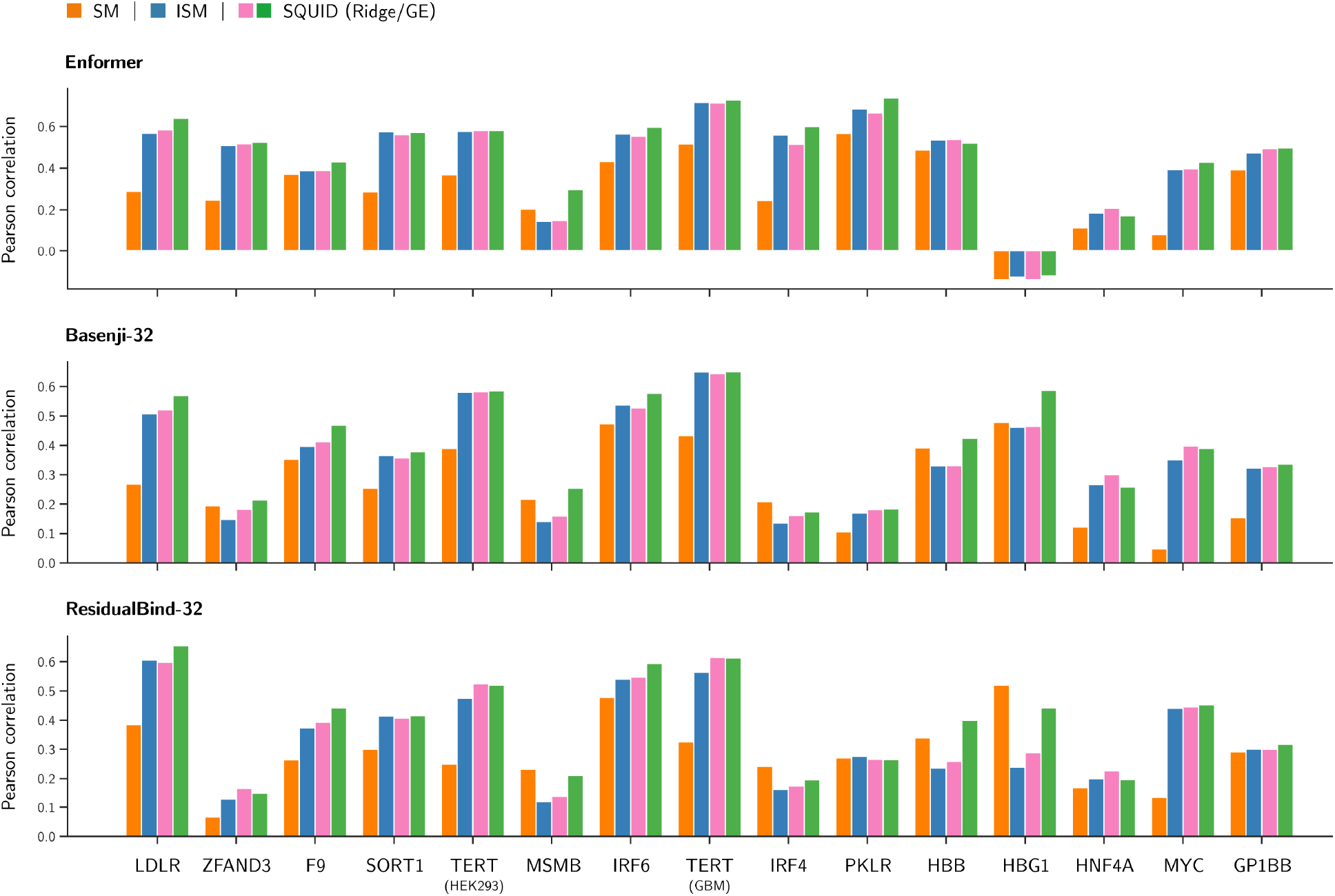
Performance of attribution methods at predicting variant effects at individual loci. Pearson correlation scores for each of the 15 disease-associated loci assayed in CAGI5, computed for the attribution methods and DNNs listed in Table 1. DNN, deep neural network.

**Supplementary Figure 2.**
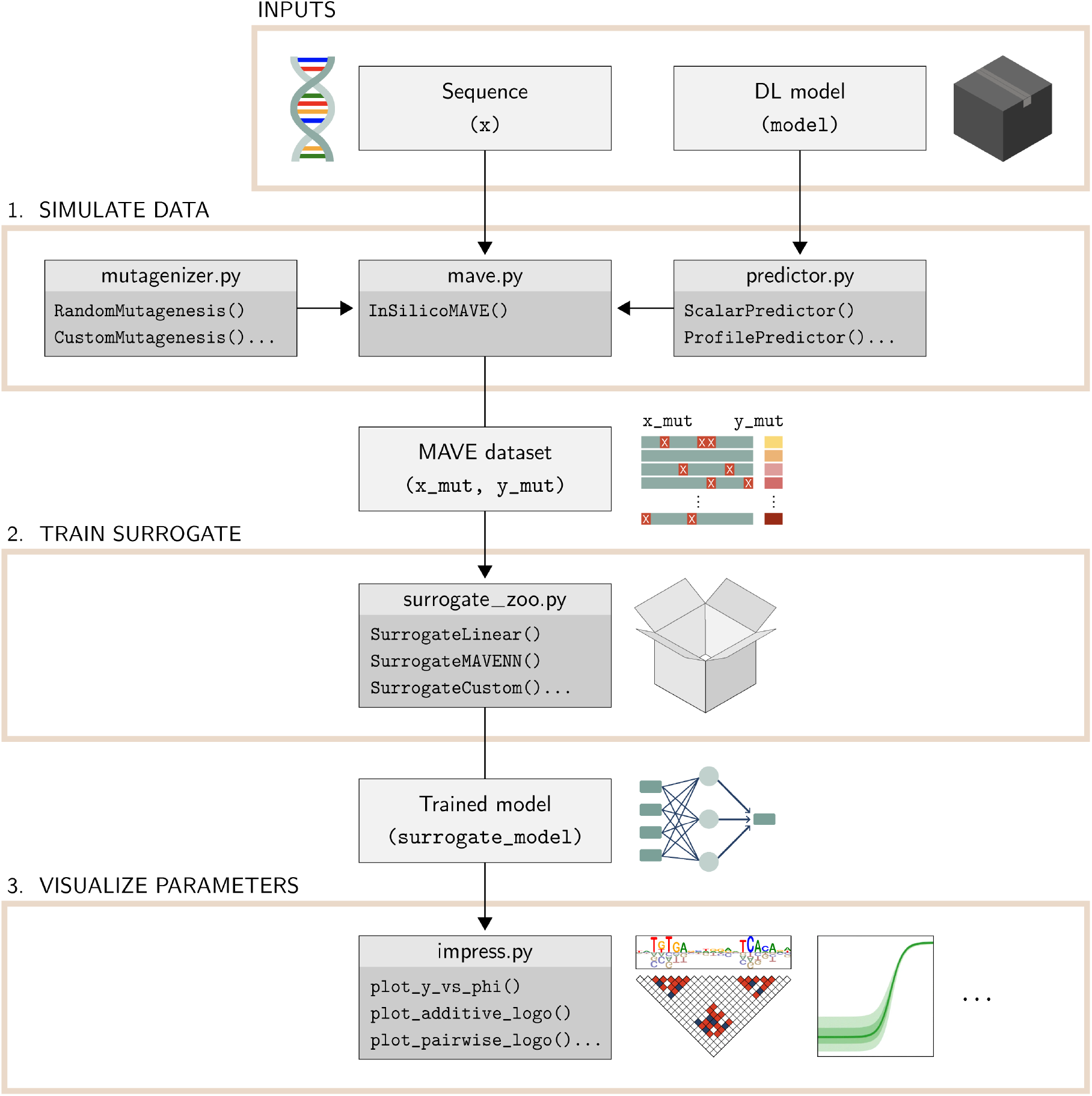
SQUID workflow. Flowchart representing a typical DNN interpretation analysis pipeline using SQUID. DNN, deep neural network.

**Supplementary Figure 3.**
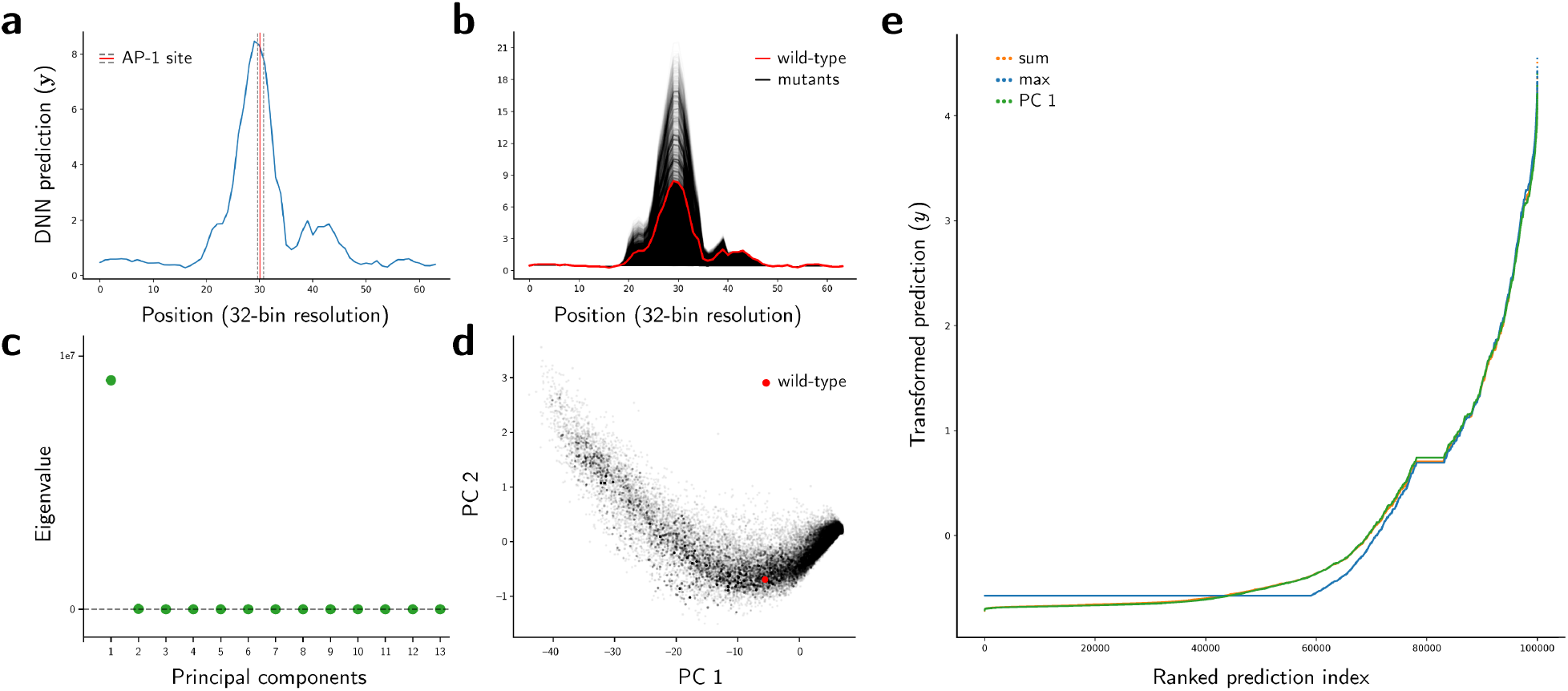
Dimensionality reduction of DNN predictions using PCA. **a**, Example ATAC-seq profile predicted by ResidualBind-32 for a representative sequence of interest containing a putative AP-1 binding site. The profile shown was cropped to a region spanning the putative site and 30 nt of flanking DNA on either side. **b**, Profiles computed for sequences in the *in silico* MAVE dataset generated by SQUID when analyzing the sequence of interest from panel **a**. **c**, Ranked eigenvalues from a PCA analysis of the profiles in panel **b**. **d**, Projection of profiles onto the first two principal components. **e**, Scalar predictions *y* for three projection methods: PCA, sum, and max. PCA projections were computed by projecting profiles onto the first principal component. Sum projections were computed by summing the entries in each profile. Max projections were computed by taking the maximum entry in each profile. To aid comparisons between different projection methods, the *y* values for each method were centered about zero and rescaled to have unit standard deviation. The flat region observed near ranked prediction index 80,000 results from sequences in the *in silico* MAVE library that have no mutations. PCA, principal component analysis.

